# Nuclear and organelle genome assemblies of five *Cucumis melo* L. accessions, Ananas, Canton, PI 414723, Vedrantais and Zhimali, belonging to diverse botanical groups

**DOI:** 10.1101/2024.10.25.620199

**Authors:** Javier Belinchon-Moreno, Aurelie Berard, Aurelie Canaguier, Isabelle Le-Clainche, Vincent Rittener-Ruff, Jacques Lagnel, Damien Hinsinger, Nathalie Boissot, Patricia Faivre-Rampant

## Abstract

The construction of accurate whole genome sequences is pivotal for characterizing the genetic diversity of plant species, identifying genes controlling important traits, or understanding their evolutionary dynamics. Here, we generated the nuclear, mitochondrial and chloroplast high-quality assemblies of five melon (*Cucumis melo* L.) accessions representing five diverse botanical groups, using the Oxford Nanopore sequencing technology. The accessions here studied included varied origins, fruit shapes, sizes, and resistance traits, providing a holistic view of melon genomic diversity.

The final chromosome-level genome assemblies ranged in size from 359 to 365 Mb, with approximately 25x coverage for four of them multiplexed in half of a PromethION flowcell, and 48x coverage for the fifth, sequenced individually in another half of a PromethION flowcell. Contigs N50 ranged from seven to 15 Mb for all the assemblies, and very long contigs reaching sizes of 20-25 Mb, almost compatible with complete chromosomes, were assembled in all the accessions. Quality assessment through BUSCO and Mercury indicated the high completeness and accuracy of the assemblies, with BUSCO values exceeding 96% for all accessions, and Mercury QV values ranging between 32 and 47.

We focused on the complex NLR resistance gene clusters to validate the accuracy of the assemblies in highly complex and repetitive regions. Through Nanopore adaptive sampling, we generated accurate targeted assemblies of these regions with a significantly higher coverage, enabling the comparison to our whole genome assemblies.

Overall, these chromosome-level assembled genomes constitute a valuable resource for research focused on melon diversity, disease resistance, evolution, and breeding applications.

**Article Summary:** This study presents high-quality nuclear, mitochondrial, and chloroplast genome assemblies for five diverse melon (*Cucumis melo* L.) accessions, using Oxford Nanopore sequencing. The assemblies represent a broad spectrum of melon diversity, including differences in origin, fruit morphology, or resistance traits. The genomes, ranging from 359 to 365 Mb, were assembled at a chromosome level with high contiguity, and verified using different validation approaches. This study provides valuable insights for research on melon genetic diversity, disease resistance, and breeding applications. The genome data will be especially valuable for plant geneticists, breeders, and researchers working on crop improvement and resistance traits.

## Introduction

Cucurbits gather numerous species that are food crops, with melon (*Cucumis melo*) being notably important. Melon originated in Asia (Sebastian, 2010) and a highly effective reproductive barrier isolated it from its relatives (Chen & Adelberg, 2020). Nevertheless, a diverse array of shapes, colors and flavors is observed in melon fruits. This large diversity is actively used for melon breeding, with more than 82 new accessions annually registered in the EU Common Catalogue (EUPVP, 2023). A recent DNA analysis of wild *C. melo* and its most important cultivar groups, revealed that modern melon cultivars go back to two lineages diverging 2 million years ago, one restricted to Asia and another to Africa (Endl et al., 2018).

As other cucurbits, melon faces many pests and diseases (Grumet et al., 2021). Plant breeders need therefore to screen a wide diversity of accessions to identify immunity loci that can be used in crop improvement programs. In plants, immunity traits are often encoded by resistance genes (R genes), commonly grouped in highly complex and diverse genome regions. Among R genes, Nucleotide-binding site leucine-rich repeat resistance genes (NLRs) constitute the largest family (Barragan & Weigel, 2021). The availability of diverse reference genomes is of great help to succeed in the identification of these genes.

Melon is a diploid species with a genome size under 400 Mb and organized in twelve chromosomes. To date, more than 40 melon genomes have been reported (Belinchon-Moreno et al., 2023), with different qualities and assembly levels, the last three being telomere-to-telomere (G. Li et al., 2023; Mo et al., 2024; Wei et al., 2023). The first assembled melon genome was released in 2012 (Garcia-Mas et al., 2012), later refined using optical mapping (Ruggieri et al., 2018) and PacBio single-molecule-real-time (SMRT) sequencing technology (Castanera et al., 2020). Additional assemblies of diverse melon lines from different origins using long read sequencing, like Payzawat (Zhang et al., 2019) and Harukei-3 (Yano et al., 2020), suggested large genomic variations, notably on chromosomes 5, 6 and 10. A recent work by Oren et al., (2022) released the assembly of 24 accessions using ONT sequencing, and revealed differences between subspecies, including size disparities and large inversions, especially on chromosomes 1 and 11.

Technological improvements, particularly long-reads sequencing technologies, have increased the accuracy of genome assemblies (Rao et al., 2021). However, some complex regions like NLR clusters remain difficult to accurately assemble, as shown in Chovelon et al., (2021) and Belinchon-Moreno et al., (2023). Targeted approaches, such as hybridization capture, Cas-9 assisted targeting methods, or Nanopore adaptive sampling, hold promise for resolving putative assembly errors (Belinchon-Moreno et al., 2023; Hook & Timp, 2023). These methods offer enhanced coverage in specific genome regions at a lower cost, facilitating more accurate genome assemblies of those areas.

Here, we report the nuclear, mitochondrial and chloroplast genome assemblies of five diverse melon genomes, belonging to five botanical groups: two belonging to the subspecies *agrestis* and three to the subspecies *melo*. We evaluated the structural diversity among the five assemblies; we focused on the NLR gene clusters to assess their accuracy in complex and repetitive genome regions. Finally, we compared our assemblies with the version already published for three of them.

## Materials and methods

### SAMPLE SELECTION

We selected five melon accessions belonging to the subspecies *melo* (Ananas, Canton and Vedrantais) and *agrestis* (PI 414723 and Zhimali). These accessions belong to five botanical groups: *ameri, reticulatus, cantalupensis, momordica* and *chinensis*, respectively; and exhibit diversity of colors and shapes (Figure 1). Their origin is located in Turkey, China (Guangdong), France, India (Uttar Pradesh) and China (Hebei), respectively. Genome assemblies of the accessions Ananas (Ananas Yoqne’am), Vedrantais and PI 414723 were already published using ONT long reads for the three of them (Oren et al., 2022) and PacBio long reads for Ananas Yoqne’am (Vaughn et al., 2022). We recovered the seeds from the INRAE Centre for Vegetable Germplasm in Avignon (Salinier et al., 2022) and cultivated in a greenhouse at INRAE GAFL, Avignon, France. We sampled young plant leaves and froze them in liquid nitrogen for subsequent DNA extraction.

**Figure 1.**
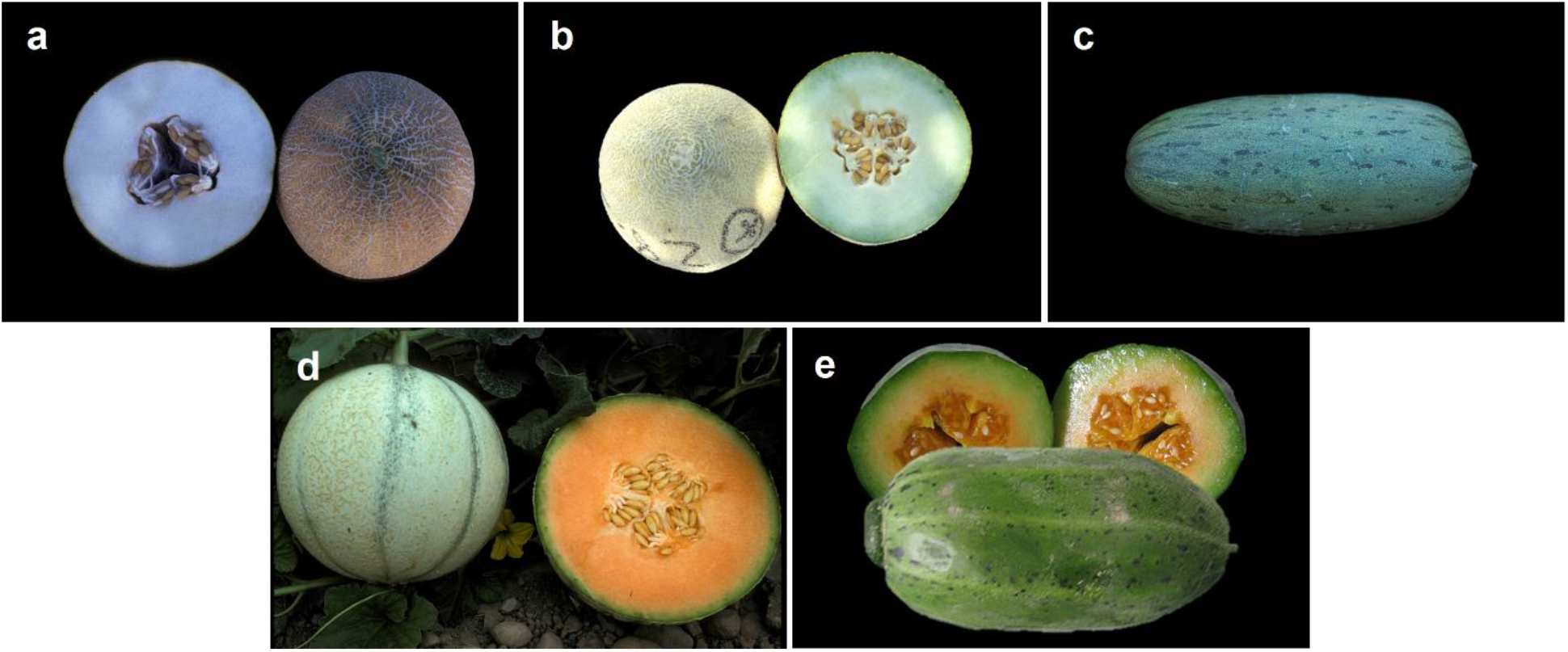
Mature fruits of the five selected accessions: a) Ananas (*ameri*); b) Canton (*reticulatus*); c) PI 414723 (*momordica*); d) Vedrantais (*cantalupensis*); e) Zhimali (*chinensis*).

### DNA EXTRACTION, LONG READS LIBRARY PREPARATION AND SEQUENCING

We extracted genomic DNA using the NucleoSpin Plant II kit (Macherey-Nagel, Germany) following the manufacturer’s protocol. We evaluated DNA quantity and quality using a Qubit4® 1x dsDNA BR Assay Kit (Invitrogen, Carlsbad, CA, USA) and an Agilent 2200 TapeStation (Agilent Technologies, Santa Clara, CA, USA). We carried out ONT libraries preparation and sequencing following ONT guidelines with the modifications described in (Belinchon-Moreno et al., 2023). We sequenced DNA of Canton using a whole PromethION *R10*.4.1 flowcell (ONT, Oxford, UK). Moreover, we multiplexed and sequenced DNAs of Ananas, PI 414723, Vedrantais and Zhimali using another PromethION *R10*.4.1 flowcell (ONT, Oxford, UK). We used barcodes NB04, NB05, NB06 and NB07 for Zhimali, PI 414723, Vedrantais and Ananas, respectively. In both experiments, we performed whole genome sequencing (WGS) using the channels 1501-3000 of the PromethION flowcells, keeping the channels 1-1500 for a Nanopore adaptive sampling (NAS) approach. We used NAS to target 15 clusters of NLR genes detected with NLGenomeSweeper v. 1.2.1, following the process established in Belinchon-Moreno et al., (2023) and using the same reference Anso77 (File S1: Table S1).

We extended both runs during 96 hours. We performed a library reloading (washing flush) in both experiences when the percentage of sequencing pores dropped to 20-30%. It corresponded to 48 hours after the start of the run in the case of Canton and 46 hours in the case of the four multiplexed samples. We set the sequencing speed to 260 bps (accuracy mode), and the quality score threshold to 10. We performed live base calling of the raw ONT FAST5 files with Guppy (ONT, London, UK) v. 6.3.9 for Canton and v. 6.4.6 for the four multiplexed samples in the “super accurate base-calling” mode. We used the MinKNOW software (ONT, London, UK) v. 22.10.7 for Canton and v. 22.12.5 for the four multiplexed DNAs. In this second run, we switched on the “trim barcodes” option to automatically trim the barcodes. For each run, we retained the “sequencing_summary.txt” file and the FASTQ files of the samples for further data processing.

### DNA EXTRACTION, SHORT READS LIBRARY PREPARATION AND SEQUENCING

For PI 414723, Vedrantais and Canton, Illumina (Illumina, San Diego, CA, USA) DNA libraries were prepared starting from DNA extracted with the Qiagen DNAeasy Plant miniKit (Qiagen, Valencia, CA, US) and using the Kapa Hyper Prep Kit PCR-Free (KapaBiosystems, Wilmington, MA, USA), following the manufacturer’s recommendations. The libraries were sequenced on an Illumina NovaSeq6000 instrument (Illumina, San Diego, CA, USA) using the 150 bp paired-end reads configuration.

For Ananas and Zhimali, MGI (MGI, Shenzhen, China) DNA libraries were prepared using the MGIEasy PCR-Free DNA Library Prep Kit (MGI, Shenzhen, China) and following manufacturer’s instructions. Briefly, 1 μg DNA was fragmented followed by a two-step size selection process using AMPure XP beads (Beckmann Coulter Genomics, Danvers, MA, USA). Quality and quantity of the fragmented and size-selected DNA was evaluated using an Agilent 2100 Bioanalyzer (Agilent Technologies, Santa Clara, CA, USA) and a Qubit4® 1x dsDNA HS Assay Kit (Invitrogen, Carlsbad, CA, USA), respectively. Then, the size-selected DNA was end-repared, 3’-adenlylated, and MGIEasy PF Adapters (MGI, Shenzhen, China) were ligated to it. The ligation products were purified using AMPure XP beads (Agilent Technologies, Santa Clara, CA, USA), and their quality and quantity was evaluated as previously described. Then, DNA was heat-denatured and the single-strand molecules were circularized. Remaining linear molecules were digested, obtaining a single-strand DNA library that was purified using AMPure XP beads (Agilent Technologies, Santa Clara, CA, USA) and quantified using a Qubit4® 1x ssDNA HS Assay Kit (Invitrogen, Carlsbad, CA, USA). Samples were pooled and DNA nano balls (DNBs) were produced and quantified with a Qubit4® 1x ssDNA HS Assay Kit (Invitrogen, Carlsbad, CA, USA). We loaded the DNBs into G400HM flowcells and sequenced them using a MGI DNBSEQ-G400RS device with a paired-end read length of 150 bp and the HotMPS High-throughput Sequencing chemistry.

### DATA PROCESSING AND GENOME ASSEMBLY

We performed data processing of ONT reads as described Belinchon-Moreno et al., (2023). Briefly, we applied two filters to the raw ONT reads for each accession. First, we filtered out reads flagged as “FAIL”, meaning that their average quality score was lower than 10. Then, using the “sequencing summary” file, we selected those reads with a “Signal Positive” end reason, filtering out those labeled as “Data Service Unblock Mux Change” (reads rejected in NAS), “Unblock Mux Change”, “Mux Change” and “Signal Negative”. For each accession, we split the reads by channel producing two different FASTQ files: one with the reads sequenced on channels 1-1500 for NAS, and another with the reads generated on channels 1501-3000 for WGS. We assessed statistics on the generated FASTQ files using seqkit stats v. 2.4.0 (Shen et al., 2016).

For each accession, we assembled the filtered ONT reads longer than 10 kb from the WGS half- flowcell using the Flye assembler v. 2.9.1 (Kolmogorov et al., 2019). We used default parameters and added the options –nano-hq and –read-error 0.03, as suggested in the software guidelines for the last ONT Kit V14 chemistry. To correct potential sequencing errors, we polished the generated contigs using the short reads with Pilon v. 1.23 (Walker et al., 2014) with defaults parameters and adding the options --minmq 30 --minqual 30. We performed a reference-guided scaffolding of the generated contigs into pseudomolecules using RagTag v. 2.1.0 (Alonge et al., 2022) with the function “scaffold”. We used the genome assembly of Harukei-3 as the reference for the nuclear and mitochondrial genomes. We filtered out the unanchored contigs shorter than 40 kb.

We assembled the plastome sequences separately through ptGAUL v. 1.0.5 (Zhou et al., 2023), using GenBank accession MT622320 (Bai et al., 2021) as a reference to select chloroplast ONT reads. ptGAUL uses Flye to generate a chloroplast assembly graph, and selects from it the different chloroplast paths. We filtered out unplaced contigs in the nuclear assembly matching the assembled chloroplast genome sequences.

Regarding NAS assemblies, filtered NAS reads longer than 5kb were assembled using SMARTdenovo v. 2018.2.19 (Liu et al., 2021), using defaults parameters and adding the “generate consensus” option. Generated contigs were filtered based on the presence of NBS domains as described in Belinchon-Moreno et al., (2023). Using NLR annotation tools like NLRAnnotator v. 2.1b or RGAugury v. 2.2 (P. Li et al., 2016; Steuernagel et al., 2020), and analysing the annotated NLR set in previously released *Cucumis melo* assemblies (Garcia-Mas et al., 2012; Mo et al., 2024; Pichot et al., 2022; Yano et al., 2020; Zhang et al., 2019), we discovered that six isolated NLR genes were missed in the reference used for adaptive sampling: three containing NBS domains, and three just containing TIR domains (File S1: Table S1). To recover these genes, we used the rejected NAS reads along with the WGS short reads to perform reference-guided assemblies using the Anso77 genome assembly as reference. We used the software Geneious Prime v. 2022.1.1 (http://www.geneious.com/) for this purpose. First, we obtained consensus sequences through the alignment of rejected NAS reads, and second, we polished the generated consensus assemblies using the WGS short reads.

We ran QUAST v. 5.0.2 (Gurevich et al., 2013) with default parameters to assess the metrics of the generated assemblies.

### REPEAT ANNOTATION

We masked and annotated repetitive elements on the WGS assemblies using a hybrid strategy combining *de novo* prediction and homology-based annotation. First, we used RepeatModeler v. 2.0.2 (Flynn et al., 2020) to build a *de novo* repeat library using each genome assembly. We used default parameters and we added the option –LTRStruct to run the LTR structural discovery pipeline (LTR_Harvest and LTR_retriever) and combined the results with those obtained with the default RepeatModeler pipeline. Then, we performed homology-based predictions of the repeat library on the genome assembly using RepeatMasker v. 4.1.2 (Smit et al., 2015) with default parameters and adding the options -a –s and –gff. This pipeline allowed to annotate and mask the genome assemblies.

### GENE PREDICTION AND FUNCTIONAL ANNOTATION

We conducted gene prediction on the assembled nuclear genomes using two distinct procedures. The first approach involved a combination of *ab initio*, homology-based, and transcriptome-based methods, integrated by EuGene v. 4.3 (Sallet et al., 2019). Annotation was performed using the eukaryote EGN-EP pipeline v. 2.0.3 (http://eugene.toulouse.inra.fr/), which encompasses various steps including probabilistic sequence model training, genome masking, computation of transcript and protein alignments, and detection of alternative splice sites. To aid in the identification of translated regions, we utilized five protein databases as evidence sources, including TAIR10, UniProtKB/Swiss-Prot, a plant subset of UniprotKB/TrEMBL, the proteome of *Cucumis melo* DHL92 v. 3.6.1, and a manually annotated subset of *Vat* proteins. Before alignment, sequences resembling those in REPBASE were excluded to ensure annotation precision. Transcriptome evidence included the meta- transcriptome used for Charmono melon genome annotation (Pichot et al., 2022), along with the CDS of manually annotated *Vat* genes (Chovelon et al., 2021). This pipeline also performs ribosomal DNA (rDNA) prediction using barrnap v. 0.9 (Seemann, 2018) in accurate mode with infernal v. 1.1.4 (Nawrocki & Eddy, 2013).

For the second approach, we employed an *ab initio* method utilizing deep-learning techniques, implemented with Helixer v. 0.3.3 (Holst et al., 2023; Stiehler et al., 2021). Annotations were restricted to the land plant lineage. The Helixer software was accessed at https://www.plabipd.de/helixer_main.html.

We searched for NLR loci using NLGenomeSweeper v. 1.2.1 (Toda et al., 2020), a tool that is able to predict NBS domains that might correspond to a complete or partial NLR gene. Based on Belinchon-Moreno et al., (2023) observations, we combined both annotations, keeping the genes annotated by Helixer in regions overlapping NBS domains, and taking the rest of the genes annotated by Eugene.

We used the EuGene approach to annotate mitochondrial assemblies. Concerning the chloroplast sequences, we annotated protein-coding, tRNA and rRNA genes using the web-based version of CHLOROBOX GeSeq (Tillich et al., 2017), available at https://chlorobox.mpimp-golm.mpg.de/geseq.html. We selected the options “Annotate plastid Inverted Repeat” and “Annotate plastid trans-spliced rps12”. We selected ARAGORN v. 1.2.38 (Laslett & Canback, 2004) with default settings for tRNA annotation. We also drew circular chloroplast maps using OGDRAW (Greiner et al., 2019).

We performed a functional annotation of all the predicted genes using EggNOG-mapper v. 2.1.12 (Cantalapiedra et al., 2021) with default settings. This tool mapped the gene sequences against the EggNOG database v. 5.0 (Huerta-Cepas et al., 2019), which provides comprehensive functional and orthologous information.

### QUALITY ASSESSMENT

We employed diverse tools and procedures to evaluate the quality and completeness of the genome assemblies. First, we identified telomeres and centromeres using the quarTeT software v. 1.1.6 (Lin et al., 2023). We used the quartet_teloexplorer.py script available at https://github.com/aaranyue/quarTeT with default parameters and setting the plant clade to search for the plant telomere sequence TTTAGGG. To identify centromeres, we used the quartet_centrominer.py script available at https://github.com/aaranyue/quarTeT with defaults settings and adding the repeat annotation and gene annotation files to improve performance.

Additionally, we determined the presence of the rDNA sequences in the ONT reads and in the assemblies by alignment of the assembled 45S rDNA sequence from the T2T Kuizilikjiz melon accession (Wei et al., 2023) and the 5S rDNA sequence from Waminal et al., (2014) (GenBank accession KF543344.1) using BLAST v. 2.15.0.

Then, we used Benchmarking Universal Single-Copy Orthologues (BUSCO) v. 5.2.2 (Simão et al., 2015) with the embryophyta_odb10 dataset, which consists of 1614 BUSCOs. This allowed us to assess the presence of conserved single-copy orthologous genes within the assemblies. Furthermore, we used Merqury v. 1.3 (Rhie et al., 2020) to estimate the consensus quality value (QV) and k-mer completeness of the assemblies using short reads. The QV serves as a log-scaled probability of error for the consensus base calls, providing insights into the accuracy of the assembled sequences.

Moreover, we used the assemblies obtained with NAS (with a higher read depth) to evaluate the accuracy of the whole-genome assemblies in complex regions of the genome usually difficult to assemble, like NLR clusters. We focused on the well-studied *Vat* region (Chovelon et al., 2021) to evaluate this accuracy, and we performed manual annotations of the *Vat* regions following Chovelon et al., (2021) and Belinchon-Moreno et al., (2023).

Finally, we compared the genome structures of the assembled accessions by aligning each chromosome sequence of each accession to its relative chromosome sequence of the other assemblies. We generated dot plots between the accessions using the online version of D-Genies (Cabanettes & Klopp, 2018) available at https://dgenies.toulouse.inra.fr/.

## Results

### DNA SEQUENCING AND GENOME ASSEMBLY

Almost 57 Gb of raw Nanopore sequences for WGS were obtained in the half PromethION flowcell of the four multiplexed samples, while almost 50 Gb were obtained in the half flowcell of Canton (Table 1). Concerning NAS sequencing, we obtained around 25 Gb in the half flowcell for the four multiplexed samples, and around 29 Gb in the half flowcell of Canton. We obtained a sequence depth of filtered ONT reads for the whole genome assemblies ranging from 24.14 to 27.72x for the four multiplexed samples. These reads presented a N50 size ranging from 24.08 to 26.88 kb. In the case of Canton, a sequence depth of 48.13x with a N50 of 18.94 kb was available for the whole genome assembly. Regarding filtered reads for the NAS assembly, we obtained a sequence depth on the target regions ranging from 138.92 to 182.41 for the four multiplexed samples, and a sequence depth of 199.66 for Canton. The N50 of these reads ranged from 21.11 to 23.69 kb for the four multiplexed samples, and was of 10.80 kb for Canton. Many raw reads were filtered out for Canton in both WGS and NAS approaches due to their poor quality or their very short size, producing a lower filtered data quantity than expected. This was probably due to technical factors such as a lower flowcell lot quality or an extra DNA fragmentation during library preparation. Figure S1 shows the higher sequence depth on the target regions in NAS compared to WGS after applying a same filter by size of 1 kb. These differences in sequence depth between the target regions are in accordance with our previous results (Belinchon-Moreno et al., 2023).

**Table 1.**
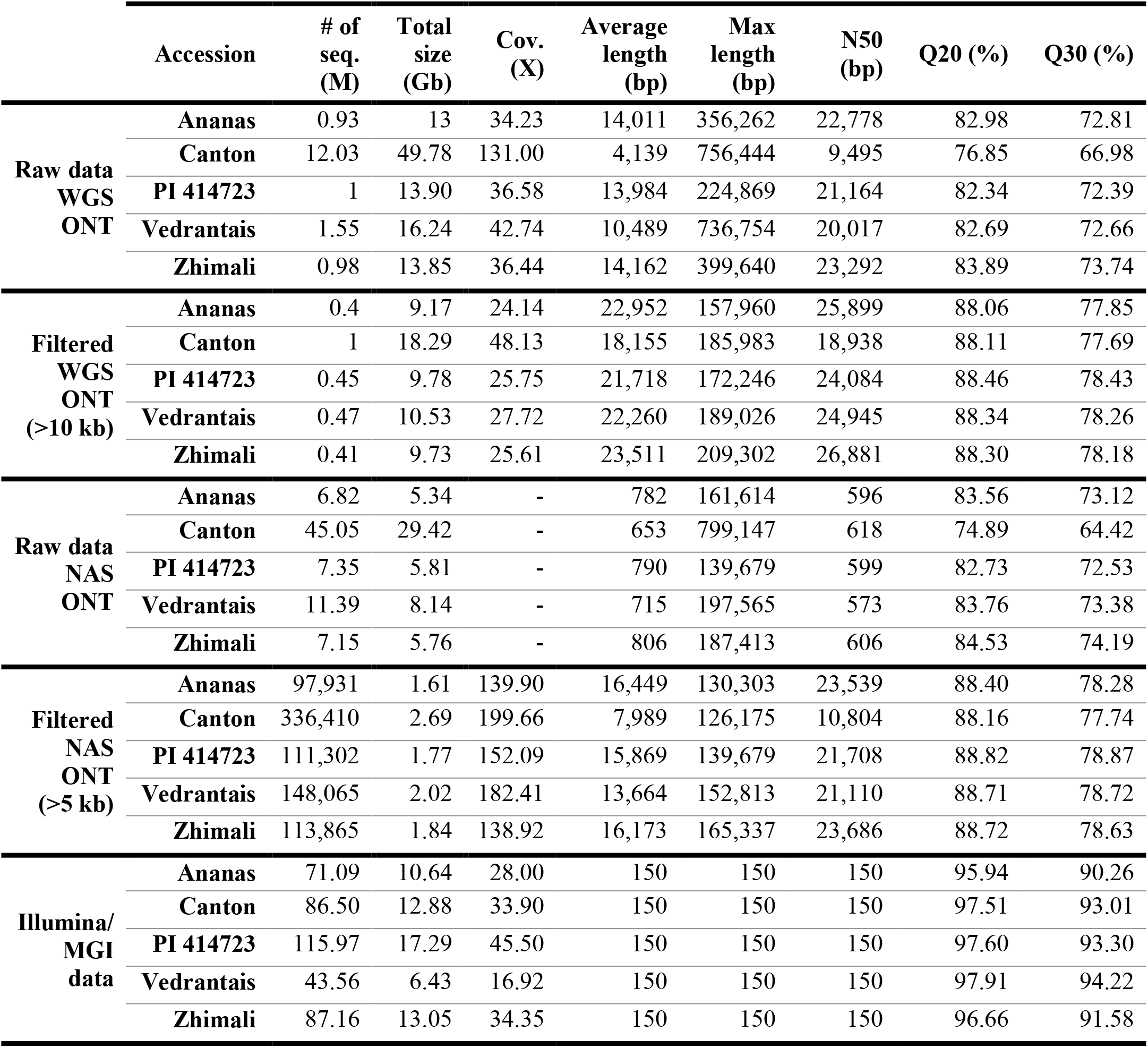
Metrics of long (WGS and NAS) and short reads for the five melon accessions.

We assembled the filtered WGS ONT reads of each accession using Flye with a subsequent polishing step with short reads using Pilon. We chose the Flye assembler due to its widely recognized superior performance with ONT sequences (Murigneux et al., 2020; Sun et al., 2021). We used BUSCO to test the importance of the polishing step by assessing genome completeness of the initial contig assembly both before and after the polishing. BUSCO were slightly better after polishing for Ananas, Canton and PI 414723, keeping stable for Vedrantais and Zhimali (File S1: Table S2), suggesting that the polishing step is not strictly necessary with the latest ONT Kit14 technology. Initial contig assemblies’ sizes ranged from 359.35 to 369.76 Mb (Table 2). Canton presented the lower number of contigs (149), while Ananas had the highest number (488). Looking at the N50 of the contigs, Ananas presented the lowest N50 (6.63 Mb), while Vedrantais showed the highest (15.12 Mb). We were able to produce some very long contigs for all accessions reaching sizes of 20-25 Mb, almost compatible with complete chromosomes. We performed the nuclear and mitochondrial anchoring of contigs using RagTag with the genome assembly of Harukei-3 as the reference. We made this choice based on the high quality of this assembly and to avoid possible assembly errors on chromosome 5 of Payzawat or 6 of DHL92, as previously established by Chovelon et al., (2021) and Oren et al., (2022). This allowed to anchor between 98.2% (for Ananas) and 99.6% (for Canton) of the total contigs size. We checked the suitability of Harukei-3 as the reference by scaffolding the *agrestis* accessions on the published *agrestis* genomes of HS (Yang et al., 2020) and PI 511890 (Mo et al., 2024). As expected, we saw no substantial differences between assemblies as our contigs sizes allow to bypass structural variations (Figure S2). After filtering out unanchored contigs shorter than 40 kb or matching their chloroplast genome sequence, from 7 (for Canton) to 36 (for Ananas) unanchored contigs were preserved in the final assemblies. The final size of the nuclear assemblies ranged from 359.21 to 365.84 Mb. Further detailed metrics of the different stages of the nuclear assemblies are summarized in Table 2.

**Table 2.**
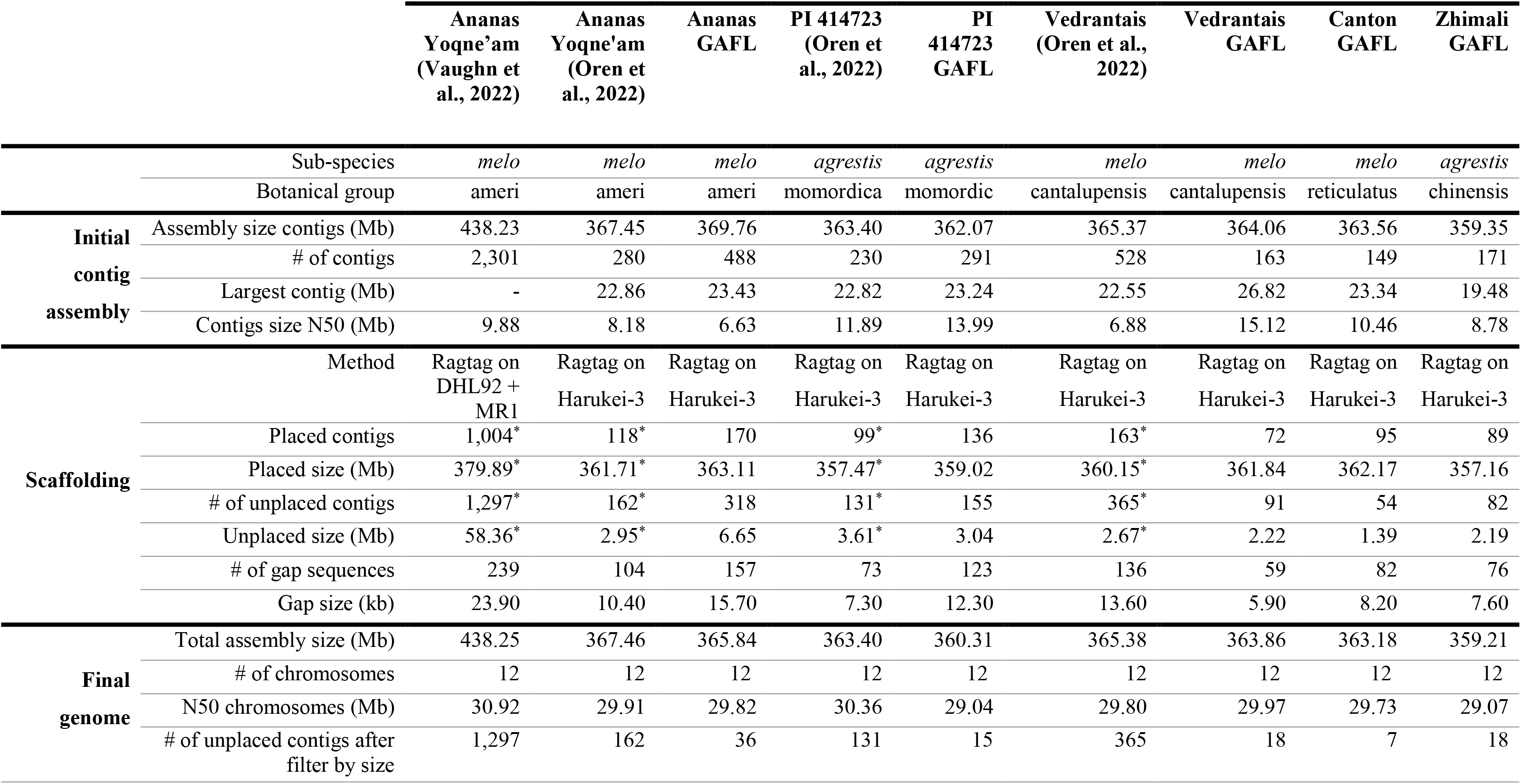
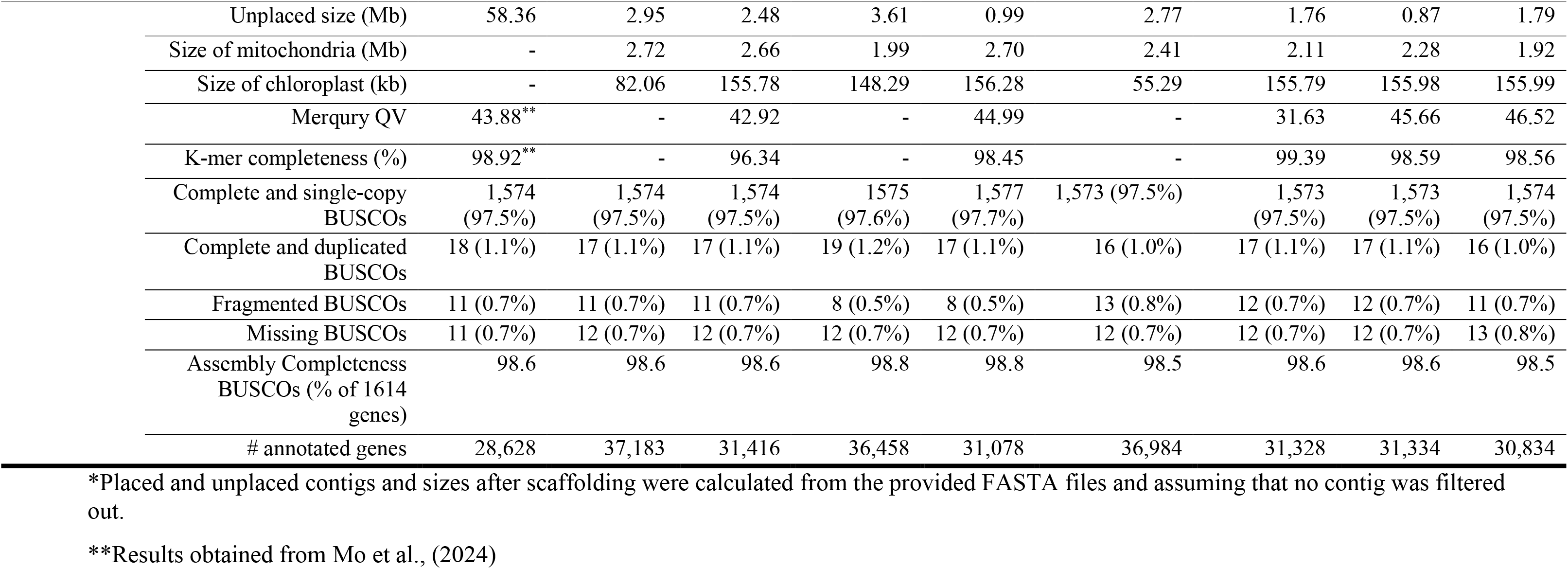
Quality metrics for various stages of the five melon whole genome assemblies. This table also includes the quality metrics for published assemblies that match the accession names of those assembled in this study.

The size of the mitochondrial sequences ranged from 1.92 Mb for Zhimali to 2.7 Mb for PI 414723 (Table 2). The chloroplast sequence was independently assembled for each accession from selected ONT reads through ptGAUL. Two paths corresponding to the two orientations of the short single-copy (SSC) were obtained for Ananas, PI 414723, Vedrantais and Zhimali, while only one path was assembled for Canton. The size of the chloroplast sequences (single path) ranged from 155.78 kb for Ananas to 156.28 kb for PI 414723 (Table 2). These sizes are similar to those found in the assembly by Wei et al., (2023).

We performed the assembly of the filtered ONT NAS reads using SMARTdenovo. As reported previously with other melon accessions (Belinchon-Moreno et al., 2023), we obtained very contiguous assemblies of the target regions with SMARTdenovo, represented each of them by one single contig in each of the assemblies. Detailed metrics of the NAS assemblies are summarized in File S1: Table S3.

### QUALITY ASSESSMENT

We used multiple strategies to evaluate the quality and completeness of the assembled genomes. First, we analyzed the assembly accuracy of tandem repetitive regions in the assemblies. These regions, such as centromeres, telomeres, and rDNA loci, are often challenging to assemble, leading to gaps in genome sequences (Wei et al., 2023). We predicted centromeric and telomeric sequences by tandem repeats and motif searching in the genome assemblies. We identified the start and end position of all 12 centromeres for all the assemblies (File S1: Table S4). Additionally, we observed 22 out of 24 telomeres for Ananas, Canton, PI 414723 and Vedrantais, and 23 for Zhimali (File S1: Table S4). All the assemblies presented sucesfully assembled telomeres at both ends in at least 10 out of the 12 chromosomes. These results show more completeness than that identified in the reference used for scaffolding (Harukei-3), where we identified a total of 14 telomeres, with 5 chromosomes presenting both of them (File S1: Table S4).

We assessed the completeness of rDNA sequences in the assemblies by alignment of the 45S and 5S rDNA sequences from Wei et al., (2023) and Waminal et al., (2014) to the ONT reads and to the assemblies. After 45S rDNA sequence alignment, we found 12 (for Canton) to 32 (for Ananas) hits with >90% identity and >5,000 bp alignment (File S1: Table S5). The number of hits on the ONT reads corrected by the coverage ranged from 22.2 (for Canton) to 59.9 (for Ananas), indicating that around half of 45S rDNA gene copies were represented in the assemblies. As reported in the assembly of Wei et al., (2023), most 45S rDNAs were located on clusters on chromosomes 4 and 10, although in Zhimali and Vedrantais over half of them were located on contigs not assigned to chromosomal location (File S1: Table S6). Moreover, we found 95 (for Vedrantais) to 222 (for Canton) hits with >90% identity and >100 bp match after 5S rDNA alignment to the assemblies (File S1: Table S5). Similarly, around 50% of the rDNA loci identified in the reads were also present in the assemblies, with the exception of PI 414723, where we found more hits in the assembly than in the ONT reads after correcting for sequence depth (File S1: Table S5). All hits across the assemblies aligned to chromosome 12, forming a 5S rDNA array with sizes ranging from 30.1 in Vedrantais to 108.6 kb in PI 414723. This 5S rDNA array was also reported in the genome assembly by Wei et al., (2023), although with a longer length of 1.8 Mb.

Then, we searched the conserved gene orthologs in the assemblies. We used the embryophyta_odb10 database for comparison purposes, as most recently published melon genome reports have employed this database (G. Li et al., 2023; Mo et al., 2024; Vaughn et al., 2022; Wei et al., 2023). From the 1614 conserved single-copy genes in the embryophyta_odb10 database, complete BUSCO scores ranged from 98.5 (for Zhimali) to 98.8% (for PI 414723) in the five assemblies. 0.5 to 0.7% orthologous genes were assigned as fragmented, and 0.7 to 0.8% were assigned as missing (Table 2). These results of complete BUSCOs are similar to those found for previous melon genomes assembled using long reads (Wei et al., 2023), including the accessions with the same name published by Oren et al., (2022) and Vaughn et al., (2022) (Table 2).

As short paired-end reads were available for the five assembled genomes, we performed a second assessment of quality and completeness using Merqury. The consensus quality value (QV) ranged from 31.63 (for Vedrantais) to 46.52 (for Zhimali), meaning that the accuracy was superior to 99.9% for all the assemblies. The k-mer completeness ranged from 96.34 (for Ananas) to 99.39% (for Vedrantais), respectively. These results were comparable to the best-assembled melon genomes published to date (Mo et al., 2024).

A last assessment of assembly quality was performed focusing on the identification of NLR resistance genes in the complete genomes and in the NAS assemblies. We searched for the conserved NBS domains using the sofwtare NLGenomeSweeper. Fifteen regions consisting on 9-10 clusters of NBS domains and 5-6 isolated NBS domains were identified in all the assemblies, in the same chromosome regions (File S1: Table S7A-E). These results agreed with those previously obtained in other melon accessions (Belinchon-Moreno et al., 2023). For Ananas, PI 414723, Vedrantais and Zhimali, the same number of NBS domains and in the same order were identified in the WGS and NAS assemblies (File S1: Table S7A,C,D,E). For Canton, two extra NBS domains were predicted in the NAS assembly: one in Region08 (where *Vat* is located), and another in Region13 (File S1: Table S7B). Further analysis of the well studied *Vat* region between markers M5 and M4 (Chovelon et al., 2021) by manual annotation of the *Vat* genes showed that the NAS and WGS assemblies of Ananas, PI 414723, Vedrantais and Zhimali were identical. We identified some small differences in the intergenic zones mostly corresponding to homopolymers. However, one extra *Vat* gene was annotated in the NAS assembly of Canton (Figure 2). An assembly error probably caused the combination of *Vat2* and *Vat3* genes into a pseudogene (*PseudoVat1*) in the WGS assembly. Long reads overpassing the concerned area demonstrated the veracity of the NAS assembly, as well as the translation of *Vat2* and *Vat3* genes into two functional *Vat* proteins. These results confirmed the high accuracy of our WGS assemblies in highly complex regions as they are NLR clusters, as almost all the predicted NBS domains were consistent between both NAS and WGS assemblies for all the accessions. In addition, the results of Canton highlighted the benefits of using NAS in complex genome regions, since its higher sequence depth output allowed to produce a better assembly, even though the WGS assembly was built with a sequence depth of 48.13X.

**Figure 2.**
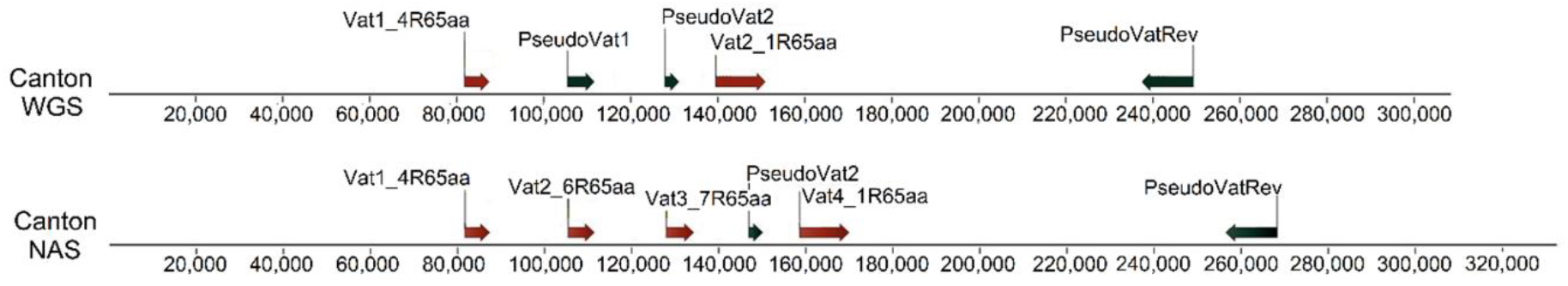
Manual annotation of the well-studied *Vat* region of Canton in the WGS (top) and NAS (bottom) assemblies.

### REPETITIVE CONTENT AND GENE ANNOTATION

The total number of annotated repetitive sequences using RepeatModeler varied from 658,089 (for PI 414723) to 674,522 (for Ananas). They represented a total size ranging from 206.42 (for PI 414723) to 211.83 Mb (for Ananas). In all the accessions, the total size of annotated repetitive sequences represented between 57 and 59 percent of the complete genome assembly size. Between 6,981 (in Zhimali) and 10,024 (in PI 414723) repetitive elements were annotated in the mitochondrial sequence, representing around 35% of the sequence length. In the chloroplast, the number of annotated repetitive elements ranged from 83 (in Zhimali) to 143 (in Vedrantais), representing around 80% of the sequence length. We compiled detailed information on the various categories of annotated repetitive elements within the nuclear, mitochondrial, and chloroplast sequences in File S1: Table S8.

We annotated protein-coding genes using a combination of EuGene and Helixer. The total number of annotated genes ranged from 30,961 for Zhimali to 31,543 for Ananas. These numbers agreed with previous published melon genome annotations (Garcia-Mas et al., 2012; Pichot et al., 2022; Yano et al., 2020; Zhang et al., 2019). From 30,638 to 31,198 gene features were annotated in the nuclear genomes. From these, 92.16 (for PI 414723) to 92.74% (for Ananas) were functionally annotated with EggNOG. The average length of gene models in the nuclear genomes ranged from 3,498 (for PI 414723) to 3,553 bp (for Vedrantais), a bit longer that previous results obtained in Payzawat (3,166 bp) and DHL92 v. 3.6.1 (2,326 bp). The average number of exons per gene in the nuclear genomes was around 4.7 for all the accessions. In total, more than 99% of the predicted protein-coding genes were allocated to the 12 chromosomes in all the assemblies. We predicted 69 (in Zhimali) to 96 (in PI 414723) genes in the mitochondrial sequences. From them, around 50% were functionally annotated in all the assemblies using EggNOG. We annotated the chloroplast genomes independently using the specialized tool CHLOROBOX GeSeq. The number of annotated genes was 127 or 128 in all the assemblies considering a single path corresponding to one orientation of the inverted repeat (IR) regions. All chloroplast assemblies exhibited the typical tetrapartite organization, with a large single copy (LSC), a small single copy (SSC), and two IR regions (Figure S3 and Figure S4). File S1: Table S9 summarizes the annotation metrics of the five genome assemblies.

### COMPARATIVE ANALYSIS

We performed chromosome-to-chromosome alignments between the five genome assemblies using D-Genies. In general, we found more co-linearity when comparing genomes from the same subspecies (*agrestis* and *melo*). Nevertheless, structural variants specific to each accession appeared in all comparisons. Large intra-chromosomal inversions located in chromosomes 1 and 11 and spanning across 1.6 and 3.2 Mb appeared when comparing the assembly of Zhimali with those of all accessions belonging to the *melo* subspecies (Figure S5 and Figure S6). This is a further evidence of the results presented by Oren et al., (2022), where they pointed out these two inversions as variants between most *agrestis* and *melo* accessions. Furthermore, as reported in the study of Oren et al., (2022), these two inversions were not found in the genome of PI 414723 (*agrestis* accession). We also found an inversion spanning across 1.6 Mb in chromosome 7 that differentiated our assembled *melo* and *agrestis* genomes (Figure S7). However, we did not find it always when checking other previously published melon assemblies. We also compared the assemblies of Ananas, PI 414723, and Vedrantais to their previously published counterparts, noting some differences. For Ananas, a 5.2 Mb fragment on chromosome 6 of the published assembly by Oren et al., (2022) was placed on chromosome 7 in ours (Figure 3a). This fragment in our assembly corresponded to an isolated contig, suggesting difficult-to-assemble regions at both ends. Comparison to the assembly of (Vaughn et al., 2022) revealed two large inversions on chromosome 6, and another on chromosome 10 (Figure 3b). The inversions on chromosome 6 likely occurred as they used the DHL92 v. 3.6.1 genome as reference for scaffolding. Regarding PI 414723, significant structural variations were also identified between the two assemblies (Figure 3c). A 2.4 Mb fragment from chromosome 2 in the published assembly was placed on chromosome 3 in ours. Additionally, a 3.1 Mb fragment at the end of chromosome 11 in the previous version was found at the beginning of chromosome 5 in our assembly. A 2.7 Mb fragment originally on chromosome 10 was relocated to chromosome 8, and a 1.2 Mb inversion was noted on chromosome 4. In the Vedrantais assemblies, three large translocations were observed (Figure 3d). A 3 Mb fragment on chromosome 2 of the previous assembly was placed on chromosome 8 in ours. Similarly, a 3.7 Mb fragment on chromosome 7 in the earlier version was found on chromosome 5 in our assembly. Additionally, a 2 Mb fragment originally on chromosome 9 was relocated to chromosome 8. For all these genomic rearrangements, the assemblies presented here were congruent with those of Harukei-3, Payzawat, DHL92, and Charmono.

**Figure 3.**
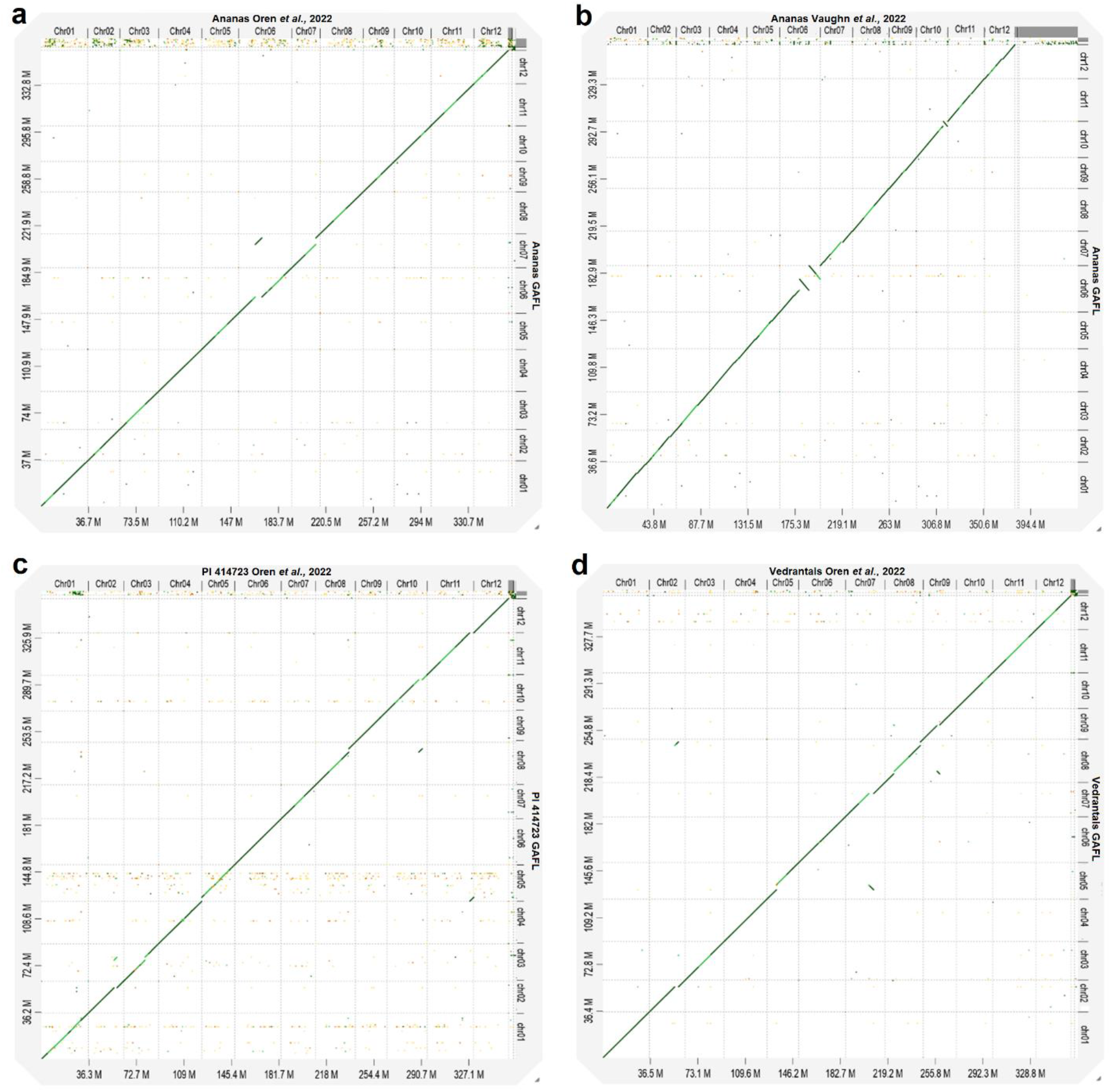
Chromosome-to-chromosome alignments between the whole genome assemblies constructed here (y-axis) and those previously published by Oren et al., (2022) or Vaughn et al., (2022) (x-axis). a) Ananas GAFL vs Ananas Oren et al., (2022). b) Ananas GAFL vs Ananas Vaughn et al., (2022). c) PI 414723 GAFL vs PI 414723 Oren et al., (2022). c) Vedrantais GAFL vs Vedrantais Oren et al., (2022).

Focusing on NLR regions, the region-to-region alignment of Ananas and Vedrantais were highly similar to those identified from the genomes presented by Oren et al., (2022) and Vaughn et al., (2022) (Figure S8a,b,d). However, significant structural variations in chromosome 1 and sequence divergence in chromosome 5 were observed between the target regions of both PI 414723 genomes (Figure S8c). By mapping the reads back to our assembly, we confirmed its accuracy. Manual annotation of the M5-M4 *Vat* zone from our assemblies and those of Oren et al., (2022) and Vaughn et al., (2022) showed identical gene structure and order for Ananas and PI 414723 accessions (Figure S9). However, one extra gene *Vat* (*Vat3*_2R65aa) was found in our assembly of Vedrantais (Figure S9). It corresponded to an insertion of 20 kb not found in the published assembly. All the differences with published assemblies might be due to assembly errors or, more likely, because both considered accessions derived after the introduction in their respective gene banks.

## Data availability

The sequencing data (ONT and short reads) and final consensus sequences of the nuclear, mitochondrial and chloroplast genomes of the five accessions are available at the NCBI database under the following BioProjects: PRJNA1164660 (Ananas), PRJNA1164662 (Canton), PRJNA1164667 (PI 414723), PRJNA1164698 (Vedrantais), PRJNA1164664 (Zhimali). Ananas: BioSample SAMN43903213, Accessions SRR30814798 and SRR30814797 for sequencing reads, and JBIEKO000000000 for genome assembly. Canton: BioSample SAMN43903216, Accessions SRR30815175 and SRR30815174 for sequencing reads, and JBHZJR000000000 for genome assembly. PI 414723: BioSample SAMN43903214, Accessions SRR30814801 and SRR30814800 for sequencing reads, and JBHZJT000000000 for genome assembly. Vedrantais: BioSample SAMN43903215, Accessions SRR30814803 and SRR30814802 for sequencing reads, and JBHZJU000000000 for genome assembly. Zhimali: BioSample SAMN43903212, Accessions SRR30806113 and SRR30806112 for sequencing reads, and JBHZJS000000000 for genome assembly. The structural and functional gene annotations of the five whole genome assemblies, as well as the repeat elements annotations are available at the Recherche Data Gouv database (https://entrepot.recherche.data.gouv.fr/) under DOI https://doi.org/10.57745/JPL1RC.

The NAS targeted sequencing data are available at the NCBI database under BioProject PRJNA1127998. Sequencing reads accessions are SRR30783378 (Ananas); SRR30783375 a (Canton); SRR30783377 (PI 414723); SRR30783376 (Vedrantais); and SRR30783379 (Zhimali). The BioSamples are the same as those specified for the WGS data. The targeted assemblies and structural gene annotations of targeted regions are available at the Recherche Data Gouv database (https://entrepot.recherche.data.gouv.fr/) under DOI https://doi.org/10.57745/ZALVPU. The large sequencing_summary.txt files used for read filtering by end reason prior assembly are available from the corresponding author on reasonable request.

## Acknowledgments

We are grateful to the Genotoul bioinformatics platform Toulouse Occitanie (Bioinfo Genotoul, https://doi.org/10.15454/1.5572369328961167E12) and the Genoscope bioinformatics platform Inti (http://dev.cng.fr/hpc/html/index.html) for providing computing and storage resources.

## Author’s contributions

J.B.M. performed the WGS and NAS experiments, the bioinformatics and statistical analyses, and all the assemblies and scaffolding. P.F.R., N.B. and D.H. conceived the study. J.L., V.C., A.C. designed and identified the ROIs for NAS. A.B. and I.L. provided expertise and experimental support. J.L. and J.B.M. performed whole genome gene annotation. V.R.R. manually annotated the *Vat* cluster for all the accessions. J.B.M., P.F.R., N.B. and D.H. wrote the manuscript. J.B.M. and A.C. did the data submission. All authors read and approved the final manuscript.

## Literature cited

1. Alonge, M., Lebeigle, L., Kirsche, M., Jenike, K., Ou, S., Aganezov, S., Wang, X., Lippman, Z. B., Schatz, M. C., & Soyk, S. (2022). Automated assembly scaffolding using RagTag elevates a new tomato system for high-throughput genome editing. Genome Biology, 23(1), 258. 10.1186/s13059-022-02823-7

2. Bai, D., Luo, X., & Yang, Y. (2021). Complete chloroplast genome sequence of ‘Field Muskmelon,’ an invasive weed to China. Mitochondrial DNA Part B, 6(12), 3352–3353. 10.1080/23802359.2021.1994888

3. Barragan, A. C., & Weigel, D. (2021). Plant NLR diversity: The known unknowns of pan-NLRomes. The Plant Cell, 33(4), 814–831. 10.1093/plcell/koaa002

4. Belinchon-Moreno, J., Berard, A., Canaguier, A., Chovelon, V., Cruaud, C., Engelen, S., Feriche- Linares, R., Le-Clainche, I., Marande, W., Rittener-Ruff, V., Lagnel, J., Hinsinger, D., Boissot, N., & Rampant, P. F. (2023). Nanopore adaptive sampling to identify the NLR-gene family in melon (Cucumis melo L.) (p. 2023.12.20.572599). bioRxiv. 10.1101/2023.12.20.572599

5. Cabanettes, F., & Klopp, C. (2018). D-GENIES: Dot plot large genomes in an interactive, efficient and simple way. PeerJ, 6, e4958. 10.7717/peerj.4958

6. Cantalapiedra, C. P., Hernández-Plaza, A., Letunic, I., Bork, P., & Huerta-Cepas, J. (2021). eggNOG- mapper v2: Functional Annotation, Orthology Assignments, and Domain Prediction at the Metagenomic Scale. Molecular Biology and Evolution, 38(12), 5825–5829. 10.1093/molbev/msab293

7. Castanera, R., Ruggieri, V., Pujol, M., Garcia-Mas, J., & Casacuberta, J. M. (2020). An Improved Melon Reference Genome With Single-Molecule Sequencing Uncovers a Recent Burst of Transposable Elements With Potential Impact on Genes. Frontiers in Plant Science, 10. https://www.frontiersin.org/articles/10.3389/fpls.2019.01815

8. Chen, J., & Adelberg, J. (2020). Interspecific hybridization in Cucumis-progress, problems, and perspectives. Horticultural Science, 35(1), 11–15.

9. Chovelon, V., Feriche-Linares, R., Barreau, G., Chadoeuf, J., Callot, C., Gautier, V., Le Paslier, M.-C., Berad, A., Faivre-Rampant, P., & Lagnel, J. (2021). Building a cluster of NLR genes conferring resistance to pests and pathogens: The story of the Vat gene cluster in cucurbits. Horticulture Research, 8.

10. Endl, J., Achigan-Dako, E. G., Pandey, A. K., Monforte, A. J., Pico, B., & Schaefer, H. (2018). Repeated domestication of melon (Cucumis melo) in Africa and Asia and a new close relative from India. American Journal of Botany, 105(10), 1662–1671. 10.1002/ajb2.1172

11. EUPVP. (2023). Common Catalogue Information System*. Available from:* https://ec.europa.eu/food/plant-variety-portal/

12. Flynn, J. M., Hubley, R., Goubert, C., Rosen, J., Clark, A. G., Feschotte, C., & Smit, A. F. (2020). RepeatModeler2 for automated genomic discovery of transposable element families. Proceedings of the National Academy of Sciences, 117(17), 9451–9457. 10.1073/pnas.1921046117

13. Garcia-Mas, J., Benjak, A., Sanseverino, W., Bourgeois, M., Mir, G., González, V. M., Hénaff, E., Câmara, F., Cozzuto, L., Lowy, E., Alioto, T., Capella-Gutiérrez, S., Blanca, J., Cañizares, J., Ziarsolo, P., Gonzalez-Ibeas, D., Rodríguez-Moreno, L., Droege, M., Du, L., … Puigdomènech, P. (2012). The genome of melon (Cucumis melo L.). Proceedings of the National Academy of Sciences of the United States of America, 109(29), 11872–11877. 10.1073/pnas.1205415109

14. Greiner, S., Lehwark, P., & Bock, R. (2019). OrganellarGenomeDRAW (OGDRAW) version 1.3.1: Expanded toolkit for the graphical visualization of organellar genomes. Nucleic Acids Research, 47(W1), W59–W64. 10.1093/nar/gkz238

15. Grumet, R., McCreight, J. D., McGregor, C., Weng, Y., Mazourek, M., Reitsma, K., Labate, J., Davis, A., & Fei, Z. (2021). Genetic Resources and Vulnerabilities of Major Cucurbit Crops. Genes, 12(8), Art. 8. 10.3390/genes12081222

16. Gurevich, A., Saveliev, V., Vyahhi, N., & Tesler, G. (2013). QUAST: Quality assessment tool for genome assemblies. Bioinformatics, 29(8), 1072–1075. 10.1093/bioinformatics/btt086

17. Holst, F., Bolger, A., Günther, C., Maß, J., Triesch, S., Kindel, F., Kiel, N., Saadat, N., Ebenhöh, O., Usadel, B., Schwacke, R., Bolger, M., Weber, A. P. M., & Denton, A. K. (2023). Helixer–de novo Prediction of Primary Eukaryotic Gene Models Combining Deep Learning and a Hidden Markov Model (p. 2023.02.06.527280). bioRxiv. 10.1101/2023.02.06.527280

18. Hook, P. W., & Timp, W. (2023). Beyond assembly: The increasing flexibility of single-molecule sequencing technology. Nature Reviews Genetics, 24(9), Art. 9. 10.1038/s41576-023-00600-1

19. Huerta-Cepas, J., Szklarczyk, D., Heller, D., Hernández-Plaza, A., Forslund, S. K., Cook, H., Mende, D. R., Letunic, I., Rattei, T., Jensen, L. J., von Mering, C., & Bork, P. (2019). eggNOG 5.0: A hierarchical, functionally and phylogenetically annotated orthology resource based on 5090 organisms and 2502 viruses. Nucleic Acids Research, 47(D1), D309–D314. 10.1093/nar/gky1085

20. Kolmogorov, M., Yuan, J., Lin, Y., & Pevzner, P. A. (2019). Assembly of long, error-prone reads using repeat graphs. Nature Biotechnology, 37(5), Art. 5. 10.1038/s41587-019-0072-8

21. Laslett, D., & Canback, B. (2004). ARAGORN, a program to detect tRNA genes and tmRNA genes in nucleotide sequences. Nucleic Acids Research, 32(1), 11–16. 10.1093/nar/gkh152

22. Li, G., Tang, L., He, Y., Xu, Y., Bendahmane, A., Garcia-Mas, J., Lin, T., & Zhao, G. (2023). The haplotype-resolved T2T reference genome highlights structural variation underlying agronomic traits of melon. Horticulture Research, 10(10), uhad182. 10.1093/hr/uhad182

23. Li, P., Quan, X., Jia, G., Xiao, J., Cloutier, S., & You, F. M. (2016). RGAugury: A pipeline for genome- wide prediction of resistance gene analogs (RGAs) in plants. BMC Genomics, 17(1), 852. 10.1186/s12864-016-3197-x

24. Lin, Y., Ye, C., Li, X., Chen, Q., Wu, Y., Zhang, F., Pan, R., Zhang, S., Chen, S., Wang, X., Cao, S., Wang, Y., Yue, Y., Liu, Y., & Yue, J. (2023). quarTeT: A telomere-to-telomere toolkit for gap-free genome assembly and centromeric repeat identification. Horticulture Research, 10(8), uhad127. 10.1093/hr/uhad127

25. Liu, H., Wu, S., Li, A., & Ruan, J. (2021). SMARTdenovo: A de novo assembler using long noisy reads. GigaByte, 2021, gigabyte15. 10.46471/gigabyte.15

26. Mo, C., Wang, H., Wei, M., Zeng, Q., Zhang, X., Fei, Z., Zhang, Y., & Kong, Q. (2024). Complete genome assembly provides a high-quality skeleton for pan-NLRome construction in melon. The Plant Journal, 118(6), 2249–2268. 10.1111/tpj.16705

27. Murigneux, V., Rai, S. K., Furtado, A., Bruxner, T. J. C., Tian, W., Harliwong, I., Wei, H., Yang, B., Ye, Q., Anderson, E., Mao, Q., Drmanac, R., Wang, O., Peters, B. A., Xu, M., Wu, P., Topp, B., Coin, L. J. M., & Henry, R. J. (2020). Comparison of long-read methods for sequencing and assembly of a plant genome. GigaScience, 9(12), giaa146. 10.1093/gigascience/giaa146

28. Nawrocki, E. P., & Eddy, S. R. (2013). Infernal 1.1: 100-fold faster RNA homology searches. Bioinformatics, 29(22), 2933–2935. 10.1093/bioinformatics/btt509

29. Oren, E., Dafna, A., Tzuri, G., Halperin, I., Isaacson, T., Elkabetz, M., Meir, A., Saar, U., Ohali, S., La, T., Romay, C., Tadmor, Y., Schaffer, A. A., Buckler, E. S., Cohen, R., Burger, J., & Gur, A. (2022). Melon pan-genome and multi-parental framework for high-resolution trait dissection (p. 2022.08.09.503186). bioRxiv. 10.1101/2022.08.09.503186

30. Pichot, C., Djari, A., Tran, J., Verdenaud, M., Marande, W., Huneau, C., Gautier, V., Latrasse, D., Arribat, S., Sommard, V., Troadec, C., Poncet, C., Bendahmane, M., Szecsi, J., Dogimont, C., Salse, J., Benhamed, M., Zouine, M., Boualem, A., & Bendahmane, A. (2022). Cantaloupe melon genome reveals 3D chromatin features and structural relationship with the ancestral cucurbitaceae karyotype. IScience, 25(1), 103696. 10.1016/j.isci.2021.103696

31. Rao, G., Zhang, J., Liu, X., Lin, C., Xin, H., Xue, L., & Wang, C. (2021). De novo assembly of a new Olea europaea genome accession using nanopore sequencing. Horticulture Research, 8, 64. 10.1038/s41438-021-00498-y

32. Rhie, A., Walenz, B. P., Koren, S., & Phillippy, A. M. (2020). Merqury: Reference-free quality, completeness, and phasing assessment for genome assemblies. Genome Biology, 21(1), 245. 10.1186/s13059-020-02134-9

33. Ruggieri, V., Alexiou, K. G., Morata, J., Argyris, J., Pujol, M., Yano, R., Nonaka, S., Ezura, H., Latrasse, D., Boualem, A., Benhamed, M., Bendahmane, A., Cigliano, R. A., Sanseverino, W., Puigdomènech, P., Casacuberta, J. M., & Garcia-Mas, J. (2018). An improved assembly and annotation of the melon (Cucumis melo L.) reference genome. Scientific Reports, 8(1), 8088. 10.1038/s41598-018-26416-2

34. Salinier, J., Lefebvre, V., Besombes, D., Burck, H., Causse, M., Daunay, M.-C., Dogimont, C., Goussopoulos, J., Gros, C., Maisonneuve, B., McLeod, L., Tobal, F., & Stevens, R. (2022). The INRAE Centre for Vegetable Germplasm: Geographically and Phenotypically Diverse Collections and Their Use in Genetics and Plant Breeding. Plants, 11(3), Art. 3. 10.3390/plants11030347

35. Sallet, E., Gouzy, J., & Schiex, T. (2019). EuGene: An Automated Integrative Gene Finder for Eukaryotes and Prokaryotes. In M. Kollmar (Ed.), Gene Prediction: Methods and Protocols (pp. 97–120). Springer. 10.1007/978-1-4939-9173-0_6

36. Sebastian, P. (2010). Cucumber (Cucumis sativus) and melon (C. melo) have numerous wild relatives in Asia and Australia, and the sister species of melon is from Australia. 10.1073/pnas.1005338107

37. Seemann, T. (2018). barrnap 0.9: Rapid ribosomal RNA prediction. *GitHub*. https://github.com/tseemann/barrnap

38. Shen, W., Le, S., Li, Y., & Hu, F. (2016). SeqKit: A Cross-Platform and Ultrafast Toolkit for FASTA/Q File Manipulation. PLOS ONE, 11(10), e0163962. 10.1371/journal.pone.0163962

39. Simão, F. A., Waterhouse, R. M., Ioannidis, P., Kriventseva, E. V., & Zdobnov, E. M. (2015). BUSCO: Assessing genome assembly and annotation completeness with single-copy orthologs. Bioinformatics, 31(19), 3210–3212. 10.1093/bioinformatics/btv351

40. Smit, A. F., Hubley, R., & Green, P. (2015). RepeatMasker Open-4.0. http://www.repeatmasker.org

41. Steuernagel, B., Witek, K., Krattinger, S. G., Ramirez-Gonzalez, R. H., Schoonbeek, H.-J., Yu, G., Baggs, E., Witek, A. I., Yadav, I., Krasileva, K. V., Jones, J. D. G., Uauy, C., Keller, B., Ridout, C. J., & Wulff, B. B. H. (2020). The NLR-Annotator Tool Enables Annotation of the Intracellular Immune Receptor Repertoire. Plant Physiology, 183(2), 468–482. 10.1104/pp.19.01273

42. Stiehler, F., Steinborn, M., Scholz, S., Dey, D., Weber, A. P. M., & Denton, A. K. (2021). Helixer: Cross-species gene annotation of large eukaryotic genomes using deep learning. Bioinformatics, 36(22– 23), 5291–5298. 10.1093/bioinformatics/btaa1044

43. Sun, J., Li, R., Chen, C., Sigwart, J. D., & Kocot, K. M. (2021). Benchmarking Oxford Nanopore read assemblers for high-quality molluscan genomes. Philosophical Transactions of the Royal Society B: Biological Sciences, 376(1825), 20200160. 10.1098/rstb.2020.0160

44. Tillich, M., Lehwark, P., Pellizzer, T., Ulbricht-Jones, E. S., Fischer, A., Bock, R., & Greiner, S. (2017). GeSeq – versatile and accurate annotation of organelle genomes. Nucleic Acids Research, 45(W1), W6– W11. 10.1093/nar/gkx391

45. Toda, N., Rustenholz, C., Baud, A., Le Paslier, M.-C., Amselem, J., Merdinoglu, D., & Faivre-Rampant, P. (2020). NLGenomeSweeper: A tool for genome-wide NBS-LRR resistance gene identification. Genes, 11(3), 333.

46. Vaughn, J. N., Branham, S. E., Abernathy, B., Hulse-Kemp, A. M., Rivers, A. R., Levi, A., & Wechter, W. P. (2022). Graph-based pangenomics maximizes genotyping density and reveals structural impacts on fungal resistance in melon. Nature Communications, 13(1), Art. 1. 10.1038/s41467-022-35621-7

47. Walker, B. J., Abeel, T., Shea, T., Priest, M., Abouelliel, A., Sakthikumar, S., Cuomo, C. A., Zeng, Q., Wortman, J., Young, S. K., & Earl, A. M. (2014). Pilon: An Integrated Tool for Comprehensive Microbial Variant Detection and Genome Assembly Improvement. PLOS ONE, 9(11), e112963. 10.1371/journal.pone.0112963

48. Waminal, N. E., Ryu, K. B., Park, B. R., & Kim, H. H. (2014). Phylogeny of Cucurbitaceae species in Korea based on 5S rDNA non-transcribed spacer. Genes & Genomics, 36(1), 57–64. 10.1007/s13258-013-0141-1

49. Wei, M., Huang, Y., Mo, C., Wang, H., Zeng, Q., Yang, W., Chen, J., Zhang, X., & Kong, Q. (2023). Telomere-to-telomere genome assembly of melon (Cucumis melo L. var. Inodorus) provides a high- quality reference for meta-QTL analysis of important traits. Horticulture Research, 10(10), uhad189. 10.1093/hr/uhad189

50. Yang, J., Deng, G., Lian, J., Garraway, J., Niu, Y., Hu, Z., Yu, J., & Zhang, M. (2020). The Chromosome-Scale Genome of Melon Dissects Genetic Architecture of Important Agronomic Traits. IScience, 23(8). 10.1016/j.isci.2020.101422

51. Yano, R., Ariizumi, T., Nonaka, S., Kawazu, Y., Zhong, S., Mueller, L., Giovannoni, J., Rose, J., & Ezura, H. (2020). Comparative genomics of muskmelon reveals a potential role for retrotransposons in the modification of gene expression. Communications Biology, 3. 10.1038/s42003-020-01172-0

52. Zhang, H., Li, X., Yu, H., Zhang, Y., Li, M., Wang, H., Wang, D., Wang, H., Fu, Q., Liu, M., Ji, C., Ma, L., Tang, J., Li, S., Miao, J., Zheng, H., & Yi, H. (2019). A High-Quality Melon Genome Assembly Provides Insights into Genetic Basis of Fruit Trait Improvement. IScience, 22, 16–27. 10.1016/j.isci.2019.10.049

53. Zhou, W., Armijos, C. E., Lee, C., Lu, R., Wang, J., Ruhlman, T. A., Jansen, R. K., Jones, A. M., & Jones, C. D. (2023). Plastid Genome Assembly Using Long-read data. Molecular Ecology Resources, 23(6), 1442–1457. 10.1111/1755-0998.13787

