## Supplemental Figures for "Nuclear and organelle genome assemblies of five *Cucumis melo* L. accessions, Ananas, Canton, PI 414723, Vedrantais and Zhimali, belonging to diverse botanical groups"

### Supplementary Figures

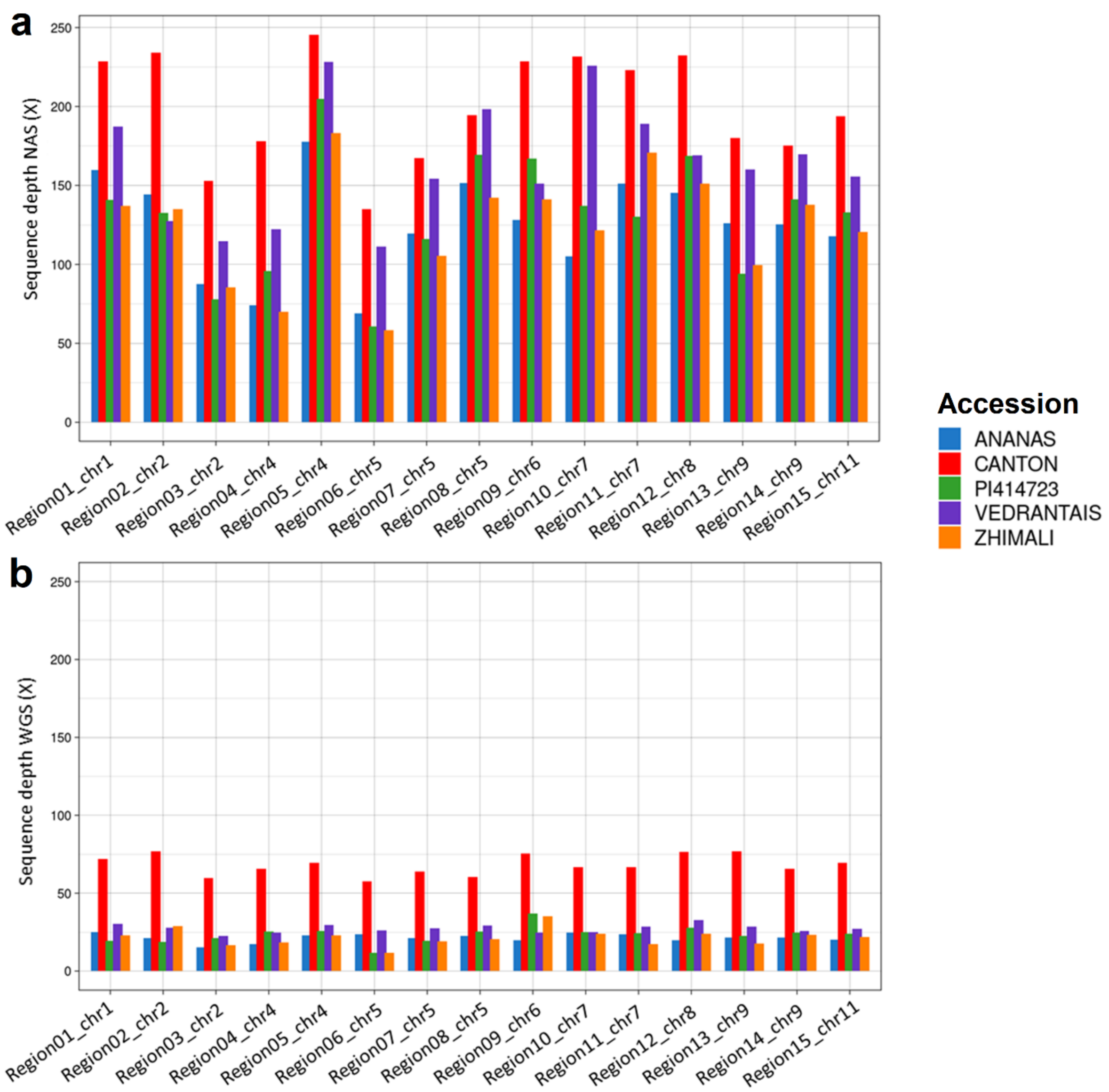


**Figure S1**. Sequence depth at the end of the run on the NAS (a) and WGS (b) half-flowcell for the five melon accessions.

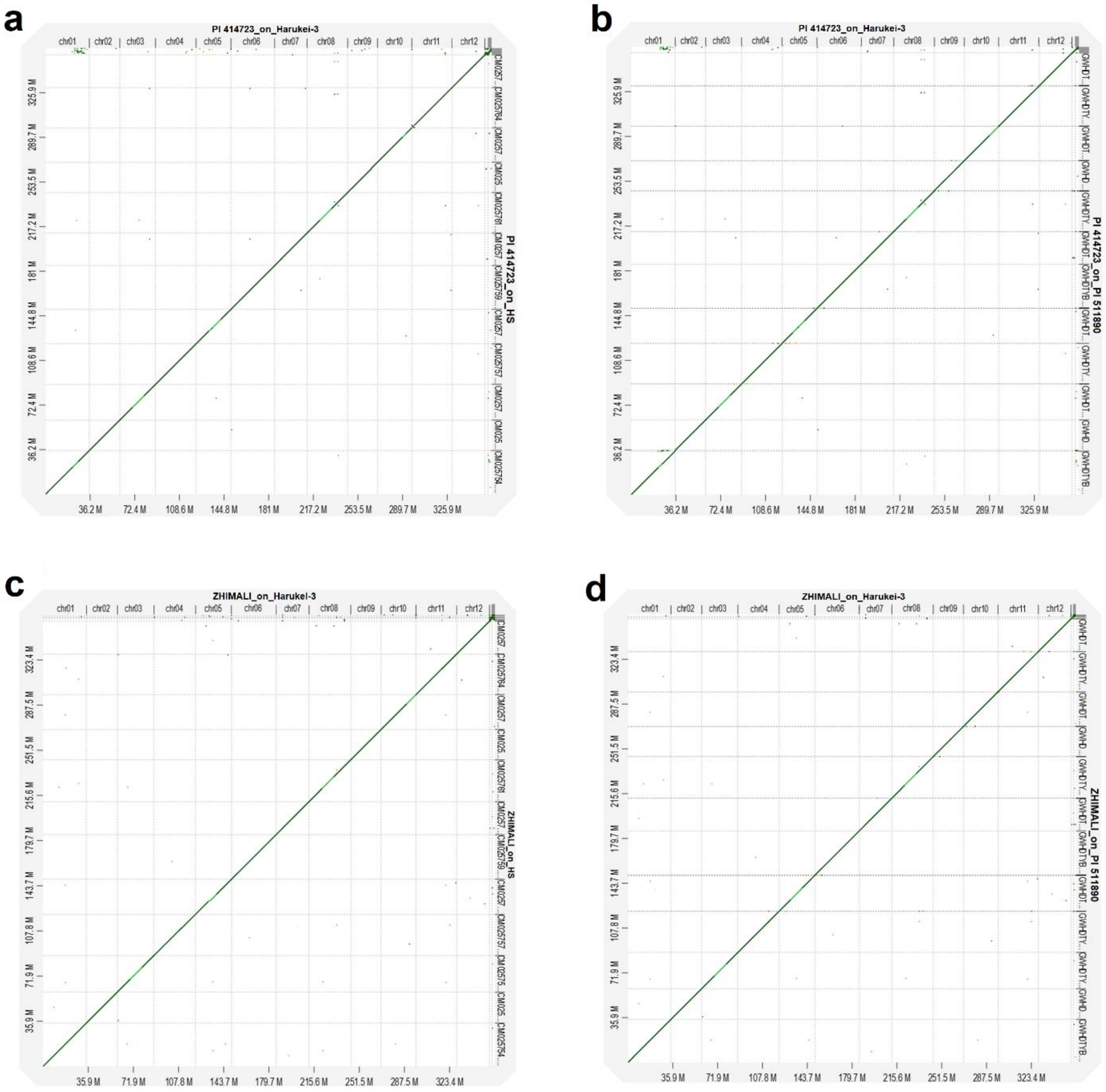


**Figure S2.** Dot plots comparing the scaffolding of *agrestis* accessions on Harukei-3, and on two published *agrestis* genome assemblies: HS and PI 511890. A) PI 414723 scaffolded on Harukei-3 (x-axis) and HS (y-axis). B) PI 414723 scaffolded on Harukei-3 (x-axis) and PI 511890 (y-axis). C) Zhimali scaffolded on Harukei-3 (x-axis) and HS (y-axis). D) Zhimali scaffolded on Harukei-3 (x-axis) and PI 511890 (y-axis).

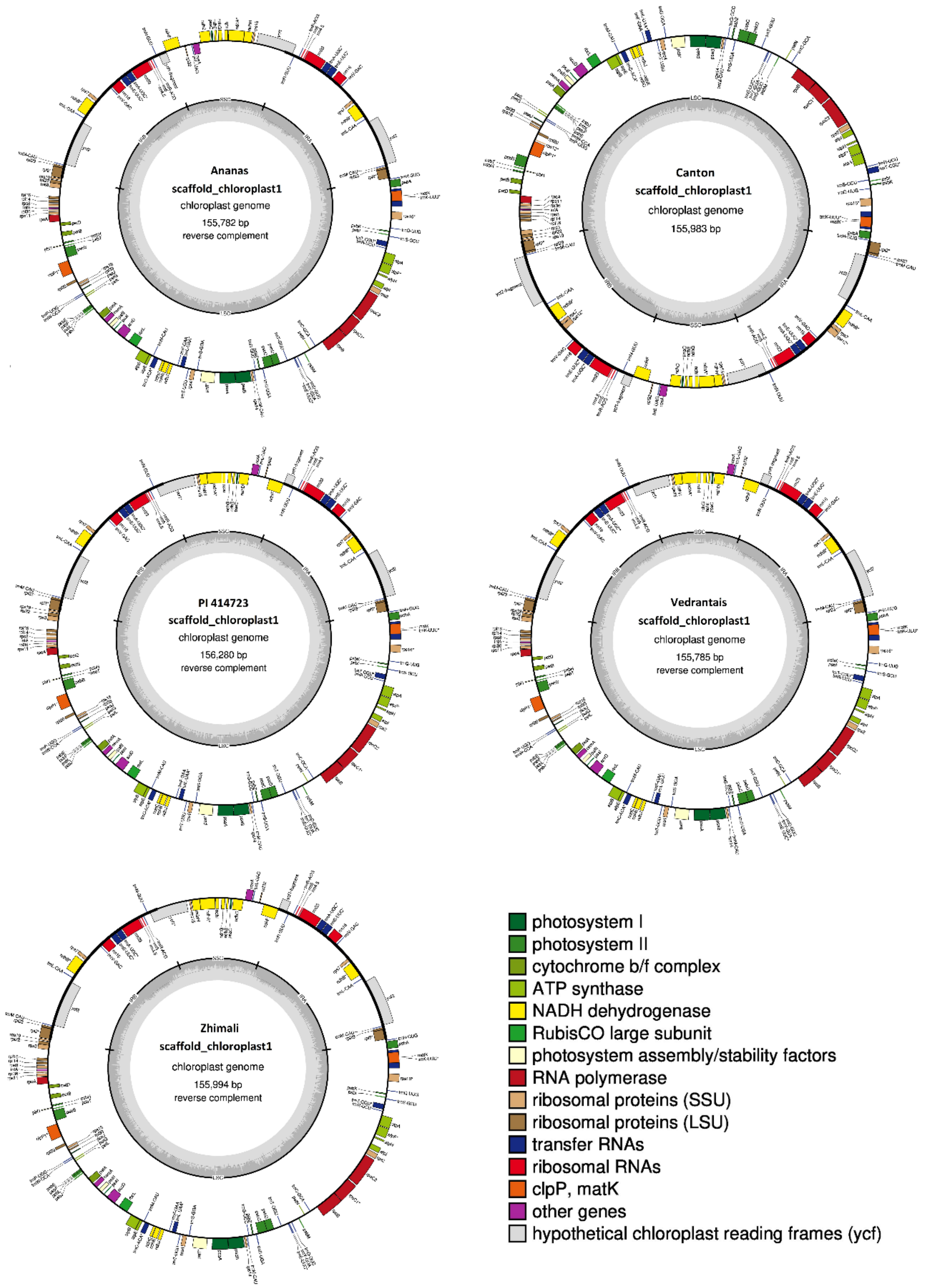


**Figure S3.** Circular diagrams depicting the first assembly path of the five chloroplast genomes, corresponding to one orientation of the SSC region. The diagram highlights the quadripartite structure, labeling the LSC, SSC, IRA, and IRB regions based on their defined boundaries. Within the inner circle, the light grey area indicates the AT content, while the darker grey layer represents the GC content. Genes situated inside the circle are transcribed in a clockwise direction, whereas those outside are transcribed counterclockwise. The genes are color-coded according to their functional groups. Genes with an asterisk denote genes containing introns.

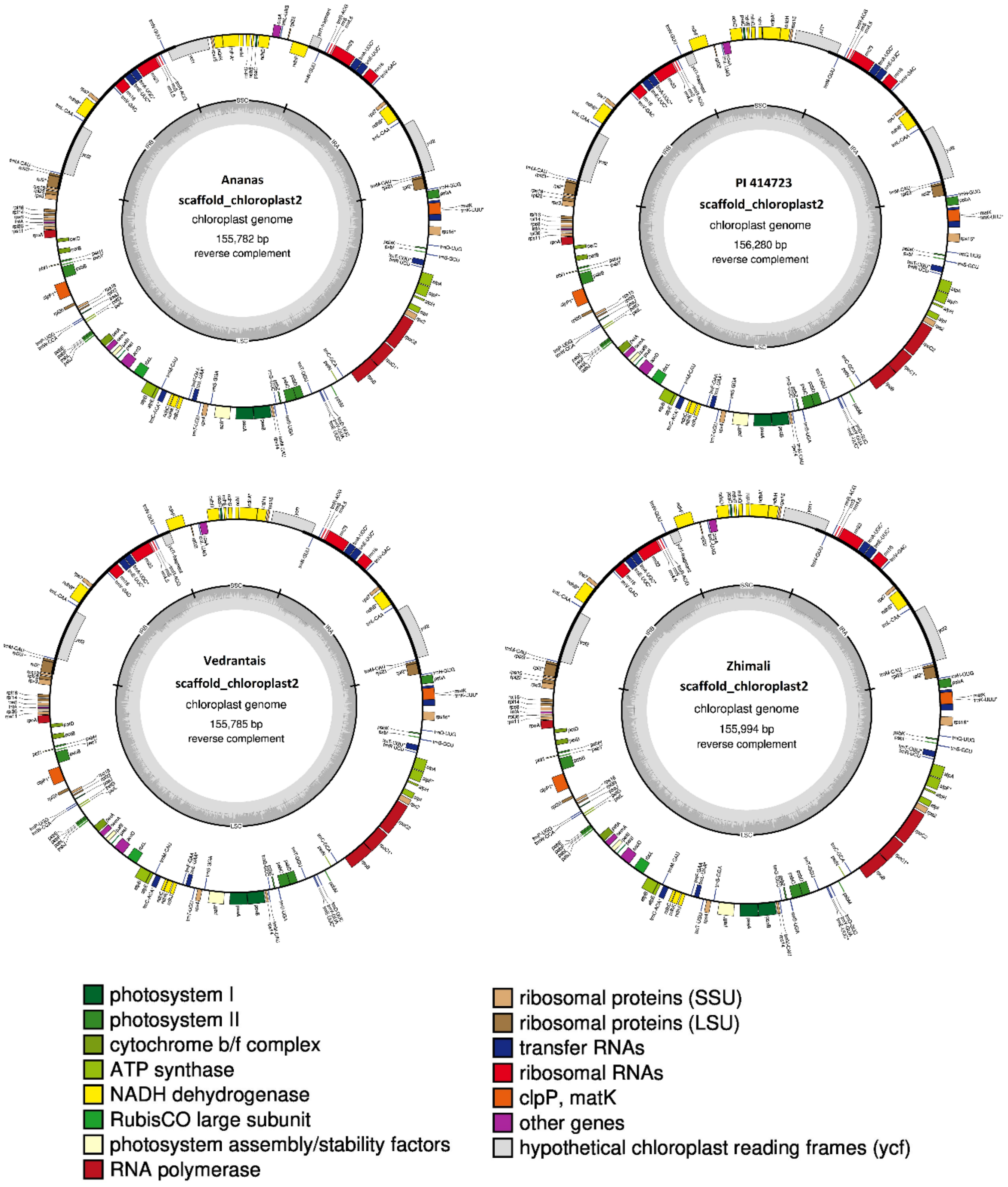


**Figure S4.** Circular diagrams depicting the second assembly path of four chloroplast genomes (Ananas, PI 414723, Vedrantais and Zhimali), corresponding to another orientation of the SSC region. The diagram highlights the quadripartite structure, labeling the LSC, SSC, IRA, and

IRB regions based on their defined boundaries. Within the inner circle, the light grey area indicates the AT content, while the darker grey layer represents the GC content. Genes situated inside the circle are transcribed in a clockwise direction, whereas those outside are transcribed counterclockwise. The genes are color-coded according to their functional groups. Genes with an asterisk denote genes containing introns.

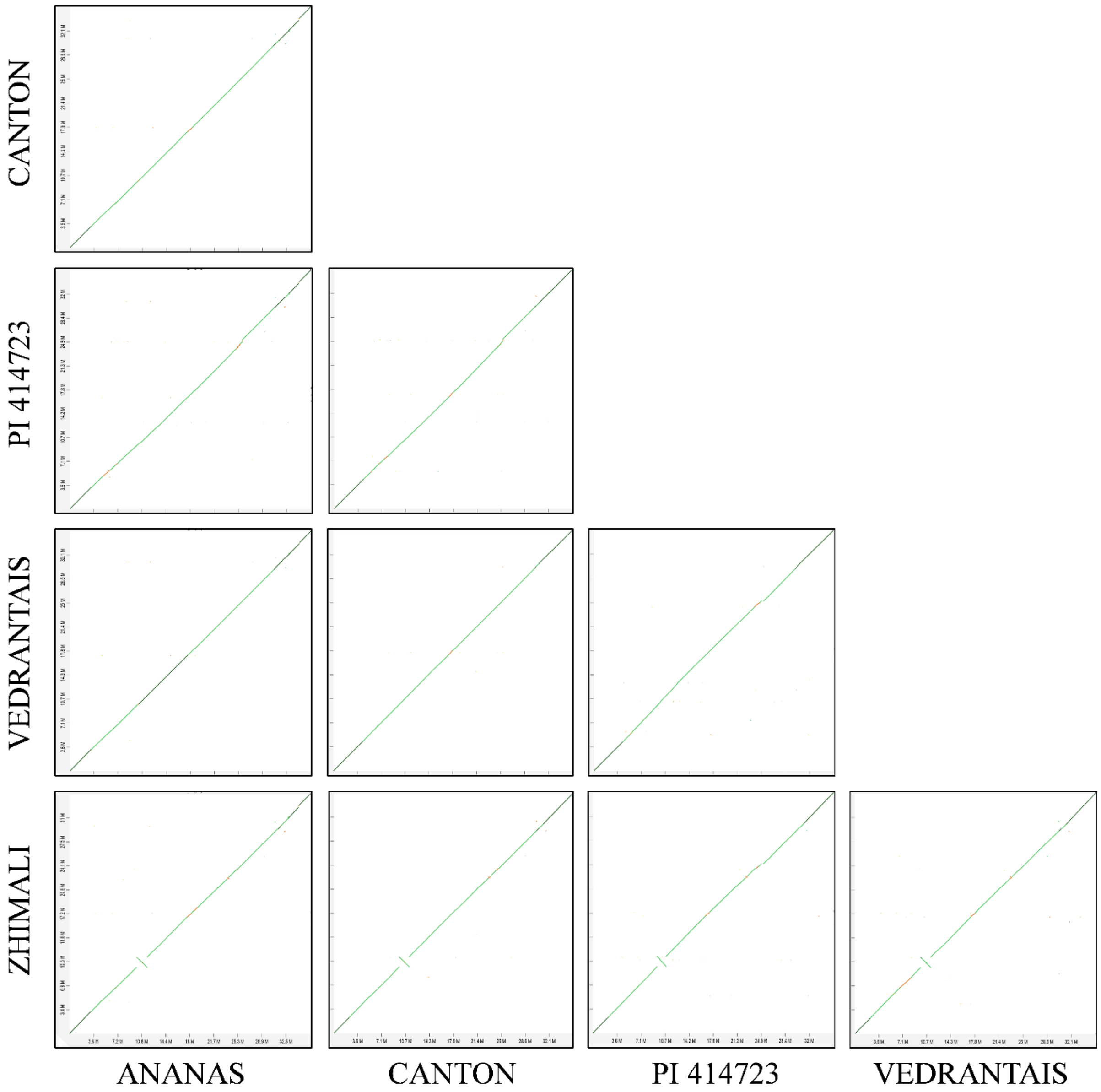


**Figure S5.** Chromosome-to-chromosome alignments between the chromosome 1 of the five genome assemblies. A large intra-chromosomal inversions spanning across 1.6 Mb appeared when comparing the assembly of Zhimali with the rest of accessions.

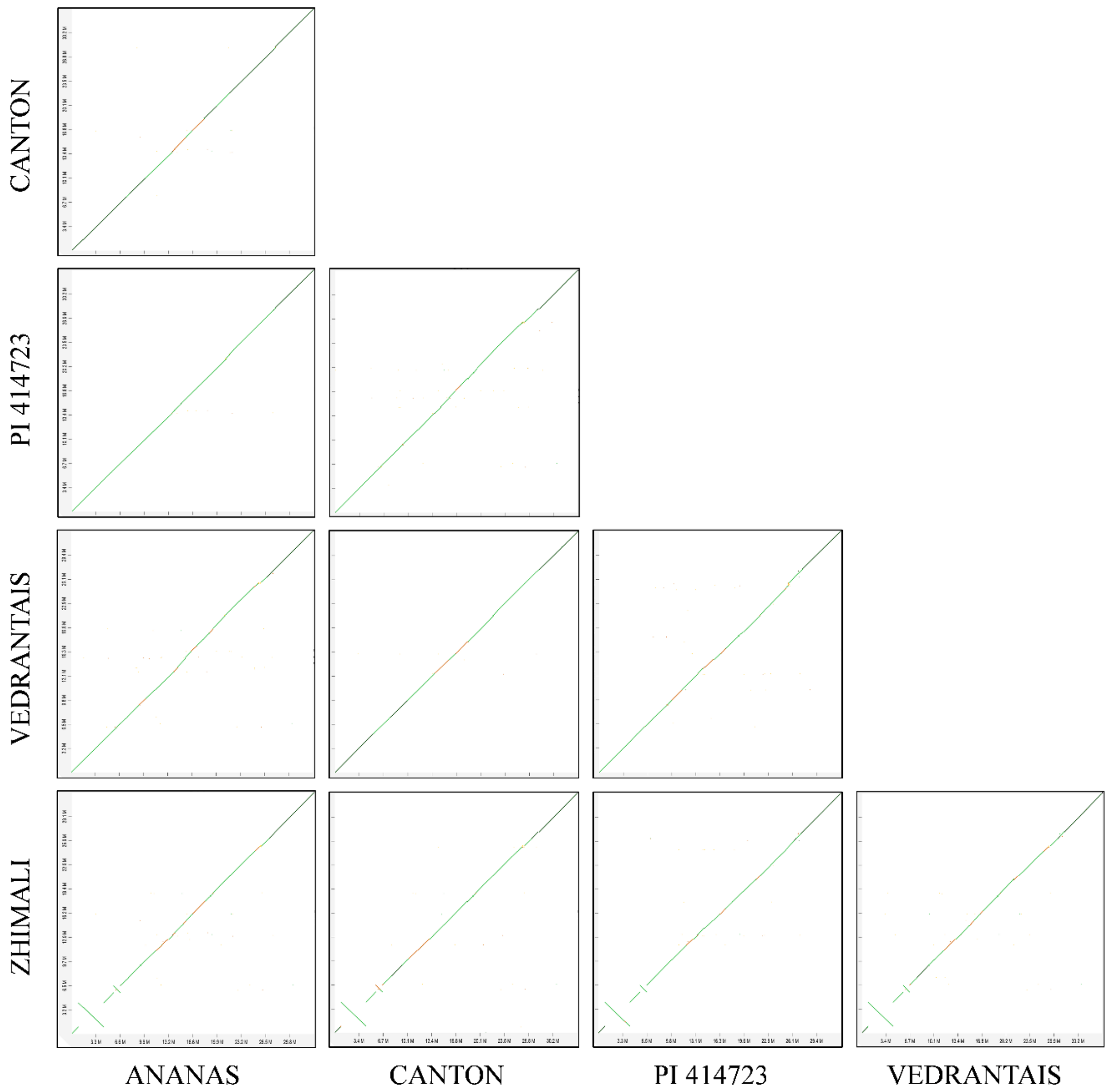


**Figure S6.** Chromosome-to-chromosome alignments between the chromosome 11 of the five genome assemblies. A large intra-chromosomal inversions spanning across 3.2 Mb appeared when comparing the assembly of Zhimali with the rest of accessions.

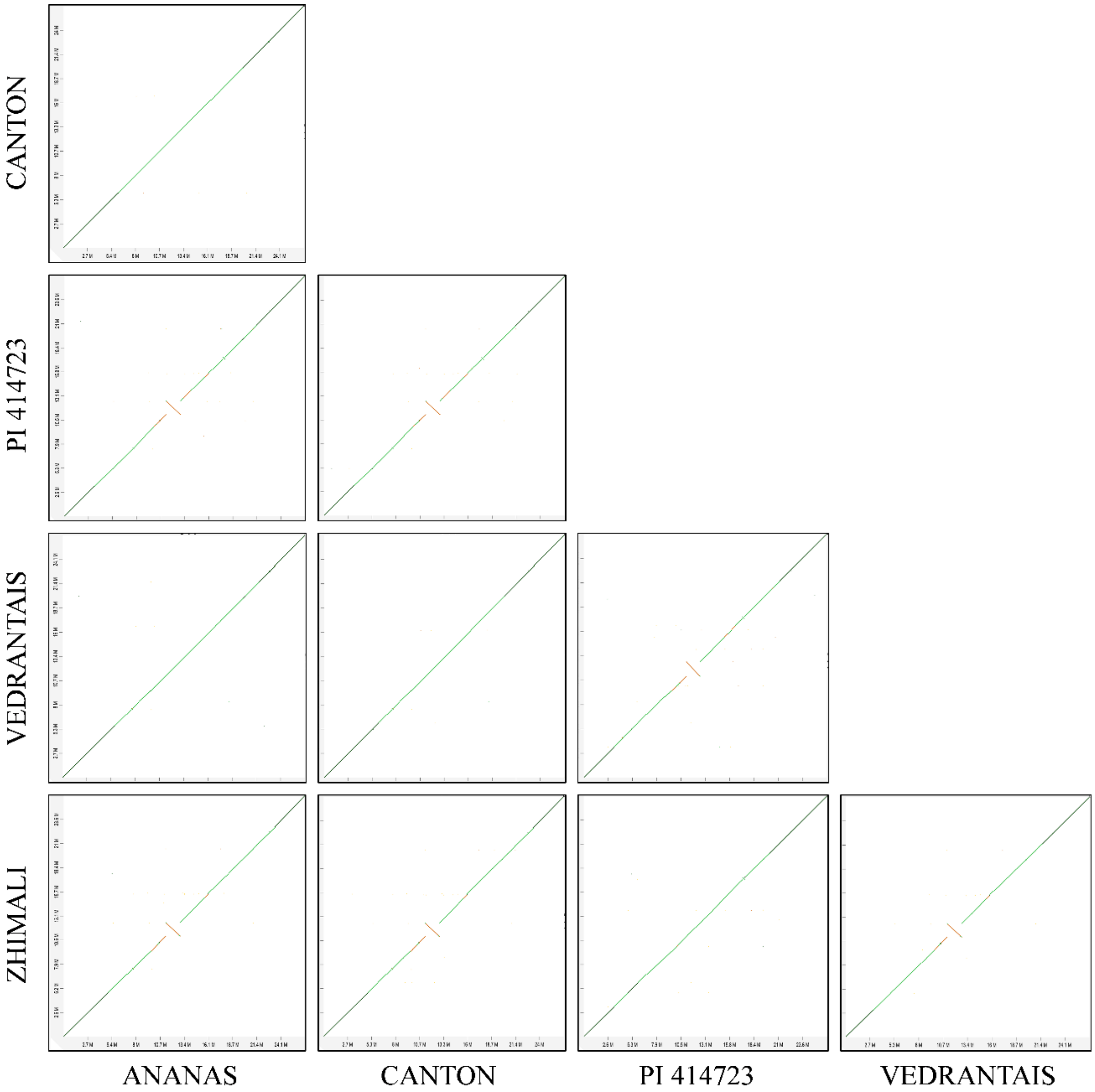


**Figure S7.** Chromosome-to-chromosome alignments between the chromosome 7 of the five genome assemblies. A large intra-chromosomal inversions spanning across 1.6 Mb appeared when comparing the assembly of *agrestis* and *melo* accessions.

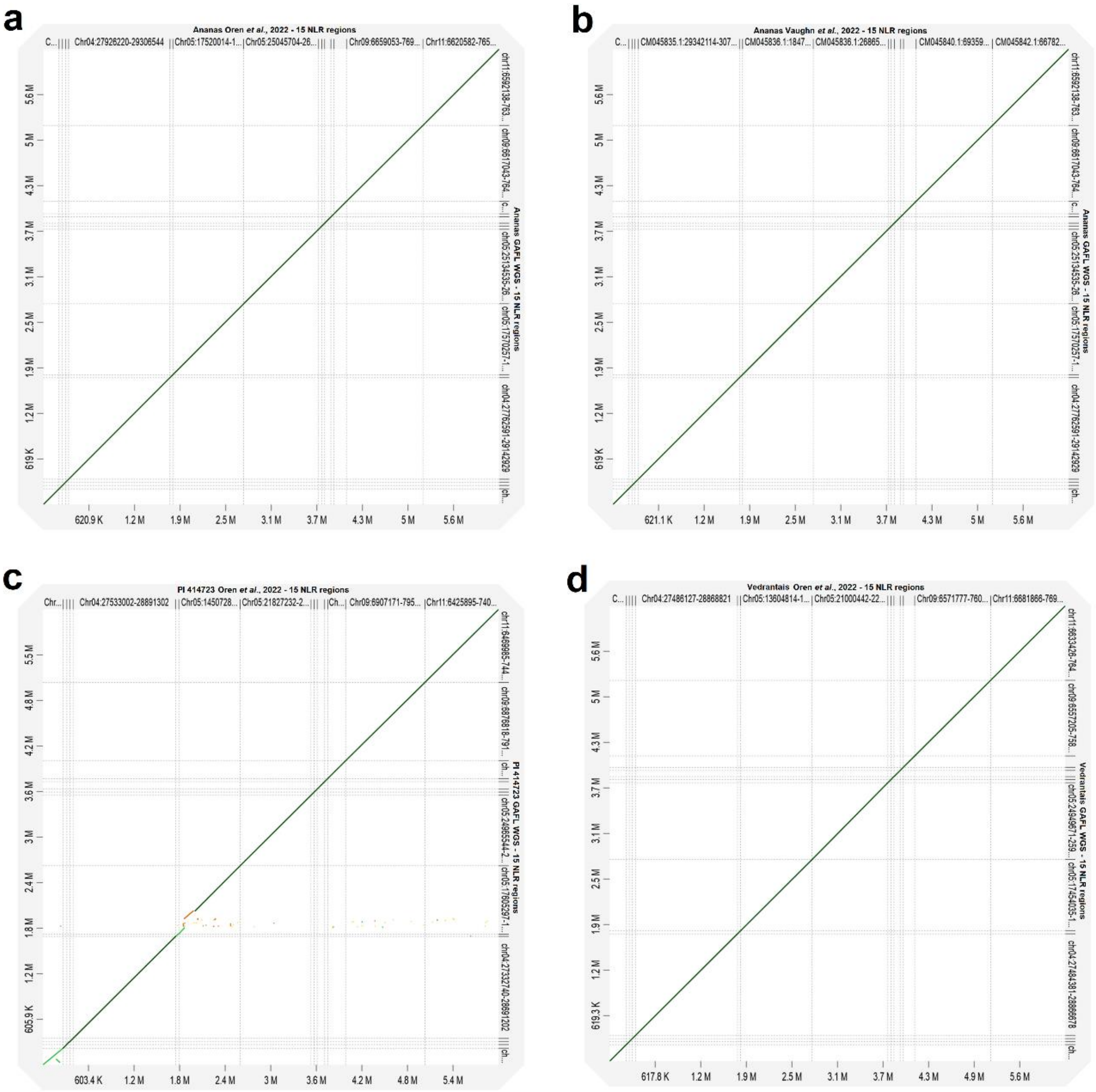


**Figure S8.** Region-to-region alignments between the 15 NLR clusters from the assemblies constructed here (y-axis) and those from the previously published genomes by Oren et al., (2022) and Vaughn et al., (2022) (x-axis). a) Ananas GAFL vs Ananas Oren et al., 2022. b) Ananas GAFL vs Ananas Vaughn et al., 2022. c) PI 414723 GAFL vs PI 414723 Oren et al., 2022. c) Vedrantais GAFL vs Vedrantais Oren et al., 2022.

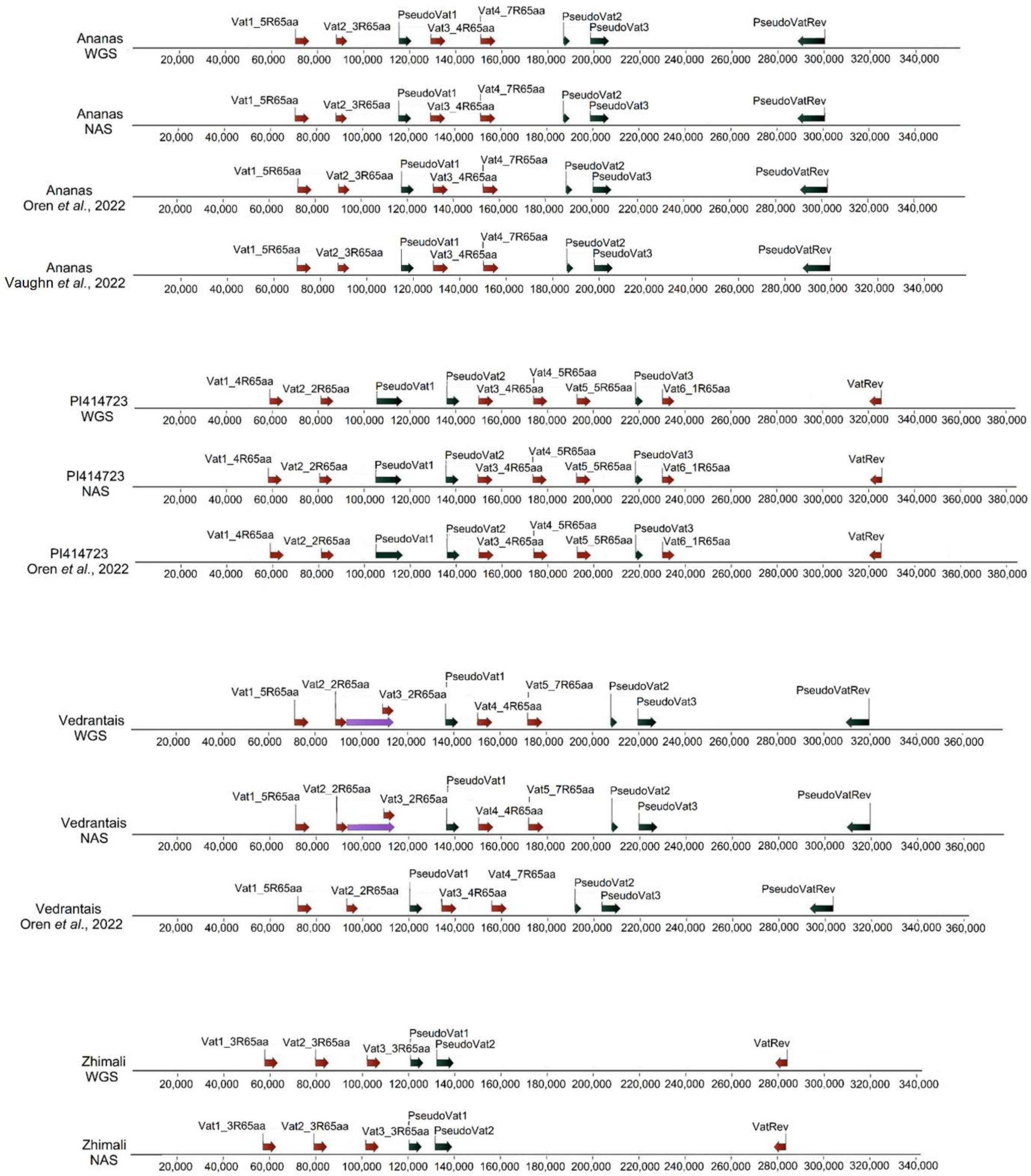


**Figure S9.** Manual annotation of the well-studied *Vat* region of Ananas, PI 414723, Vedrantais and Zhimali. We show the manual annotation of the WGS and NAS assemblies constructed here, as well as those previously published by Oren et al., (2022) and Vaughn et al., (2022) when they were available. For Vedrantais, the purple arrow represents an insertion found in both WGS and NAS assemblies compared to the previously published assembly.
